# AGAPE (computAtional G-quadruplex Affinitiy PrEdiction): The first Artificial Intelligence workflow for G-quadruplex binding affinity prediction

**DOI:** 10.1101/2024.11.14.623389

**Authors:** Luisa D’Anna, Salvatore Contino, Rosalinda Marinello, Julie Fares, Giada De Simone, Antonio Monari, Florent Barbault, Giampaolo Barone, Alessio Terenzi, Ugo Perricone

## Abstract

AGAPE (computational G-quadruplex Affinity Prediction) is a novel machine learning (ML)-based tool designed to predict the binding and stabilizing potential of small molecules targeting G-quadruplexes (G4s). G4s, prevalent in telomeres and oncogene promoters, are promising therapeutic targets, but designing selective binders remains challenging. Building upon a curated dataset of 1217 compounds labelled through Förster Resonance Energy Transfer (FRET) melting assays, AGAPE integrates 5666 molecular descriptors, both classical and quantum chemical. It captures features relevant to G4 recognition, driving researcher to predict the potential G4 stabilization of small molecules, including both organic ligands and metal complexes. Among the trained ML models, XGBoost achieved the best performance with an accuracy of nearly 91%, using 489 selected features. SHAP analysis highlighted descriptors related to molecular topology, polarizability, and electrostatic potential as key contributors to the classification. AGAPE is deployed through a user-friendly web interface supporting batch prediction and secure data handling and provides a robust and interpretable tool to accelerate the discovery of G4-stabilizing compounds, integrating quantum chemical information within an ML-driven cheminformatics framework.

## Introduction

Guanine quadruplexes (G4s) are nucleic acid structures that play critical roles in diverse biological processes[1]. They form in guanine-rich DNA and RNA sequences through Hoogsteen hydrogen bonding, resulting in stacked guanine tetrads stabilized by monovalent cations (e.g., Na^+^, K^+^, Figure 1) [2]. Despite their rigid core, G4s exhibit remarkable structural polymorphism, with variations in strand orientation, loop features, and glycosidic bond orientations leading to parallel, antiparallel, and hybrid topologies. These polymorphic features influence G4 stability and biological function [1, 3]. G4 motifs are highly conserved in telomeres and oncogene promoters, where they regulate gene expression, replication, and chromosomal stability [4, 5]. Alterations in G4 stability have been linked to diseases such as cancer and neurodegenerative disorders, underscoring their potential as therapeutic targets [6]. Small molecules designed to stabilize G4 structures typically feature extended aromatic scaffolds to promote stacking interactions with external G-quartets, as well as positively charged groups to enhance electrostatic interactions with the nucleic acid backbone[7]. However, additional features are necessary to deploy stable interactions with the G4 loops, grooves, and extruded bases [3]. Various organic and metal-based G4 binders have shown promising affinity and selectivity [8–10]. Among them, Salphen metal complexes have attracted significant interest due to their strong and selective binding to G4 structures.

**Fig. 1.**
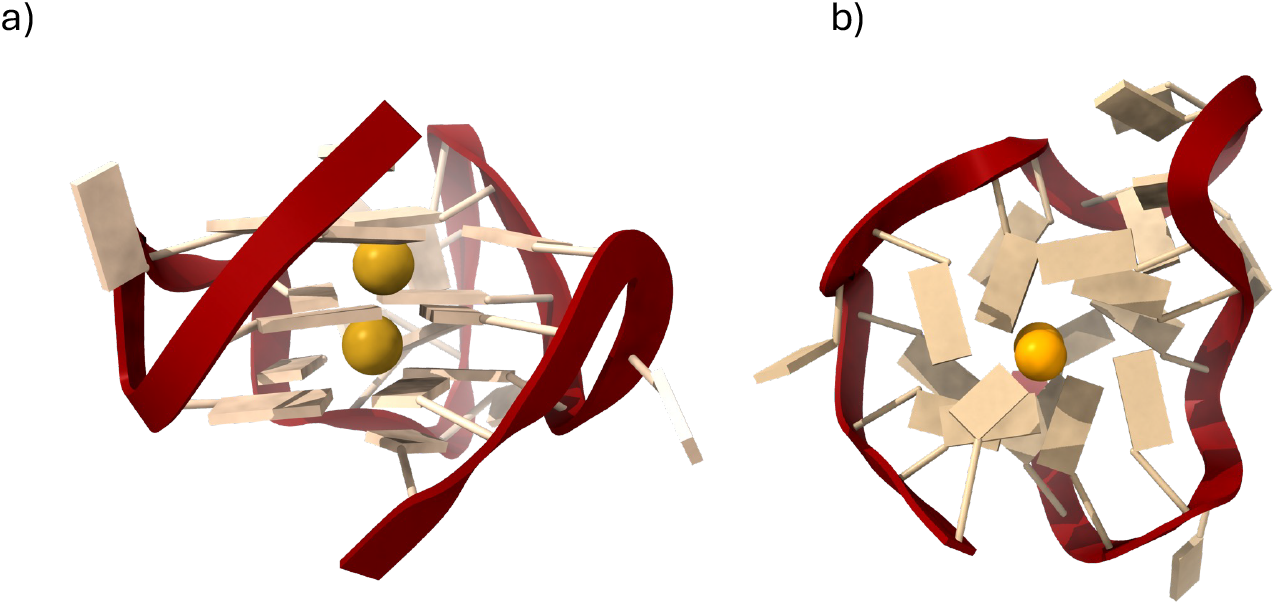
Side (a) and top (b) view of a typical parallel G-quadruplex motif.

Many transition metal complexes involving Salphen-like ligands have been investigated to elucidate the role of the central metal ion and lateral substituents in optimizing their interactions with G4 structures [10–12]. However, challenges persist in achieving high specificity for disease-relevant G4 structures. Ideal G4 binders should selectively recognize specific G4 motifs implicated in disease progression, modulating only the associated biological pathways.

The selective targeting of G4s requires a deeper understanding of the structural features that govern G4-binder interactions. Current approaches often rely on molecular modelling, quantum chemical calculations, and experimental techniques such as calorimetry and spectroscopy. However, these methods are time-consuming and resource-intensive. Conversely, machine learning (ML) offers a transformative approach, enabling the rapid analysis of large datasets to predict molecular interactions and guide drug discovery [13–15]. ML has already demonstrated success in predicting biological activities, physicochemical properties[16–19], and protein structures, as evidenced by the 2024 Nobel Prize in Chemistry awarded for advancements in computational protein design [20].

Despite the growing application of AI in drug discovery, its use in designing selective G4 binders remains unexplored. Existing AI tools for G4 research primarily focus on predicting G4 folding and stability rather than host-guest binding [21–26]. It is worth noting that the current dataset of G4 binders is relatively limited compared to other biologically relevant targets, such as kinase inhibitors [27, 28]. Still, this relative scarcity of data and specialized tools underscores the urgent need of collecting and organizing information that can contribute to building and feeding an initial data-driven framework.

To address this gap, we developed AGAPE (computational G-quadruplex Affinity Prediction), the first ML-based framework designed to predict the G4 stabilization potential of small molecules. Beyond its predictive capabilities, AGAPE aims to elucidate the key chemical features that underpin G4 binding by using explainable AI techniques to enhance model interpretability. Our approach utilizes molecular descriptors, including molecular embeddings [29, 30], to construct a binary classification model that categorizes the compounds as ACTIVE or INACTIVE based on their G4 stabilization potential.

The use of molecular descriptors in ML applications is highly valuable across various domains [31–33], as it facilitates the development of interpretable models compared to the use of molecular fingerprints for structure representation [29, 31, 32]. Given the complex molecular and electronic nature of certain G4 binders, such as metal complexes, we have also incorporated quantum chemical (QC) properties as additional features to characterize these molecules. QC molecular descriptors provide detailed insights into molecular interactions, and their integration with ML has been proven useful and efficient in predicting physicochemical properties [34–37], reactivity, and [38] regioselectivity of substitution [39], as well in the exploration of the chemical space [40]. Moreover, combining QC descriptors with ML techniques shows significant promise in medicinal chemistry for drug design, lead discovery, and optimization[41]. However, and despite its potential, this approach has not yet become a standard practice in drug design, likely due to the computational overhead associated with QC calculations. By addressing these challenges, AGAPE represents a novel and accessible tool for the chemoinformatics community, facilitating the discovery and optimization of selective G4 binders.

## Methods

### Data collection and dataset creation

Data were mainly retrieved from the public database G4LDB (v. 2.2)[42], a collection aimed to explore molecules targeting all kinds of G-quadruplexes (DNA and RNA), as well as from compounds from literature not included in G4LDB complemented by our in-house compounds library. To classify molecules in our dataset as ACTIVE or INACTIVE, we used results from Förster Resonance Energy Transfer (FRET) melting assays, using thresholds derived from benchmark compounds in the G4LDB database and supporting literature data [42, 43]. Specifically, we labelled molecules presenting Δ*T* ≤ 8 °C as “INACTIVE” and those with Δ*T* ≥ 15 °C as “ACTIVE” based on our evaluation of literature data and on the method itself [44–47]. Although a ΔT of approximately 15 °C is generally accepted in the literature as an indicator of activity, we chose a lower threshold to more reliably identify INACTIVE compounds and minimize the risk of false positives. The QC geometry optimization of the selected molecules was performed using Jaguar tool from Schrodinger suite (Schrodinger Release 2023-2: Jaguar, Schrodinger, LLC, New York, NY, 2024). Density Functional Theory (DFT) calculations, performed with B3LYP as the exchange correlation functional and the LANLD2Z basis set, was used to optimize the molecular geometry and calculate all the relevant Quantum Chemistry (QC) properties. DFT calculations are indeed particularly useful to obtain the geometry of small molecules, especially transition metal complexes. In fact, the parameterization of classical force fields for transition metal complexes is still not straightforward and force field parameters for some of the metals in our dataset are either missing or not standardized. In addition, resorting to QC approach allows to explicitly include features directly related to the electronic structure of the studied molecule without relying on any empirical assumption. Thus, we calculated QC molecular properties and retained them as additional features for our model creation. The selected QC descriptors were mainly focused on the determination of point charges distribution, polarizability, electrostatic potentials, and surface size. Starting from the QC optimized geometries we have also calculated classical molecular descriptors using the alvaDesc Software [48, 49].

### Data cleaning and preparation

Dataset has been processed into 3 consecutive steps, namely data cleaning, data transformation and dimensionality reduction.

Data Cleaning: The dataset was organized into columns representing individual variables. Columns with constant values, entirely missing values, or more than 20% of missing values were removed to improve data quality.

Data Transformation: Missing values in numeric columns containing integer have been replaced with the median value, while those in columns containing double-precision floating data points were substituted with the mean value.

Dimensionality Reduction: We calculated the Pearson correlation coefficient 53 for each pair of columns, as a measure of the correlation between the two variables. Variables with a high degree of correlation (threshold set at 0.9) were identified and subsequently removed, allowing to significantly reduce redundancy in the dataset.

After these three steps of cleaning and preparation, the residual dataset consisted of 1723 features, which were then used as descriptors for building our ML model.

### Preprocessing

The features were normalized to the range [0, 1] using the MinMaxScaler estimator. This scaler transforms each feature individually to fit within the specified range, using a linear transformation. Note that MinMaxScaler does not mitigate the effect of outliers. To prevent overfitting, two validation strategies were adopted:

- A split of 80% training, 10% validation, and 10% test set;
- 10-fold Cross Validation.

### Feature selection algorithm

Feature selection methods were used to reduce the number of properties related to G-quadruplex binding properties. Furthermore, the dimensionality reduction of the dataset allows the use of a smaller and more interpretable dataset. The main categories of feature selection used in this work are: Filter, Embedded, and Wrapper.

The Filter methodology was applied first. This approach evaluates each feature independently from the predictive model, filtering out the most irrelevant variables. It relies on statistical metrics that compute a score between each independent variable and the target. As a result, features that are redundant with respect to the target variable are excluded.

The main advantage of the filter methods is their computational efficiency. However, they do not account for interactions between features.

To reduce the dimensionality of the dataset and highlight the most relevant variables for prediction, three filter techniques were applied:

- Mutual Information: measures the statistical dependence between each feature and the target.
- ANOVA (Analysis of Variance): evaluates the statistical significance of differences between groups.
- Chi-square: measures the statistical independence between categorical variables.

The practical implementation of these filter methods was performed using the SelectKBest class from the Scikit-learn library. This class allows the automatic selection of the top *k* features based on the scores computed by the specified statistical scoring function, i.e. mutual information classification for Mutual Information, f classification for ANOVA, and chi2 for the Chi-square test.

The Embedded approach integrates feature selection within the model training process. The classifier learns which features are relevant during the training phase. This technique offers a good trade-off between computational cost and predictive performance.

Based on these results, Random Forest coupled with filter methods appeared to slightly outperform the other methods and has shown better performance stability independently from the filter method. Thus, we have also decided to evaluate the embedded feature selection capability of Random Forest by employing the feature importance measure provided by the ensemble module of Scikit-learn.

The Wrapper methodology explores combinations of features through a search strategy that considers the predictive power of subsets. While it captures feature interactions, it is computationally more expensive.

Specifically, we used a Sequential Forward Selection, which belongs to the category of “greedy” approaches. These strategies are used to reduce the initial d-dimensionality to a *k* −*dimensional* feature space where *k < d*. These approaches are based on the automatic selection of the *k* subset inherent to the identified task. This step aims to improve computational efficiency by reducing the generalisation error of the model through the removal of irrelevant or noisy features.

Starting from a set of features *Y* = {*y*_1_, *y*_2_ …,*y*_*d*_} the algorithm is initialised with an empty set ∅ so that *k* = 0 (where k is the size of the subset). The next phase involves the gradual inclusion of features as follows (equation 1)

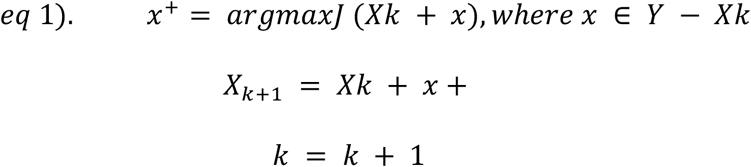

where *x*^+^ is the feature that maximises our criterion function, i.e. the feature associated with the best performance of the classifier when added to *X*_*k*_.

The procedure continues until the criterion is satisfied or until *k* = *p*, i.e. the a priori defined maximum size of subset K. In our work we have set a maximum *p* = 1723, i.e. the total number of features present in the initial dataset, to assure exploration of all the features’ space.

### Machine learning models

The following machine learning algorithms were developed and trained: Decision Tree (DT), Random Forest (RF), Gaussian Naive Bayes (NB), Support Vector Machine (SVM), and XGBoost.

DT is a supervised learning method used for classification. The goal is to create a model that predicts the value of a target variable by learning simple decision rules inferred from the data features. It is a model that offers advantages in terms of interpretation and simplicity.

Building upon the strategy exploited with decision trees, we extended our models to RF, a more advanced approach that aggregates the predictions of multiple decision trees to produce a more accurate and stable output. As an ensemble learning method, RF constructs a large number of trees during the training phase [50]. Each tree performs repeated feature splitting until a specific condition is met, using the following splitting criteria (equation 3)

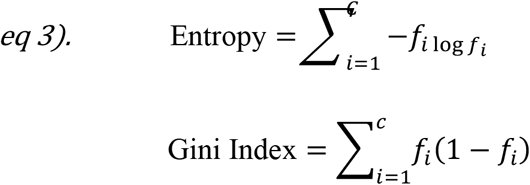

where **C** is the number of unique labels and *f*_*i*_ the frequency of label *i* at the node.

Naive Bayes (NB) is an algorithm based on Bayes theorem that assumes that all the features are mutually independent. Specifically, Gaussian NB [44] has been used, which is based on computing a normal distribution assuming that each likelihood (*P*(*x*_*i*_|*y*)) follows a normal distribution for each *x*_*i*_ with *y*, as expressed in the equation 4).

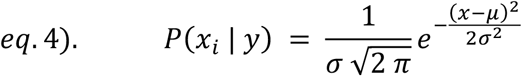

SVM is one of the most widely used and effective algorithms for classification and regression. It is based on the definition of hyperplanes to separate data into perfect groups. It is originally designed to separate classes that are easily dividable using linear kernels, but can be extended to more complex data adopting non-linear kernels. This strategy is called kernel trick and aims to maximise classification capacity [51]. Its linear application is shown in the equation 5)

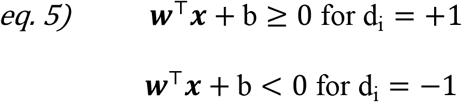

where *w* is a weight vector, *x* is the input vector and *b* is the bias.

To improve the performance, we also tested the kernel functions reported in Table 1.

**Table 1.**
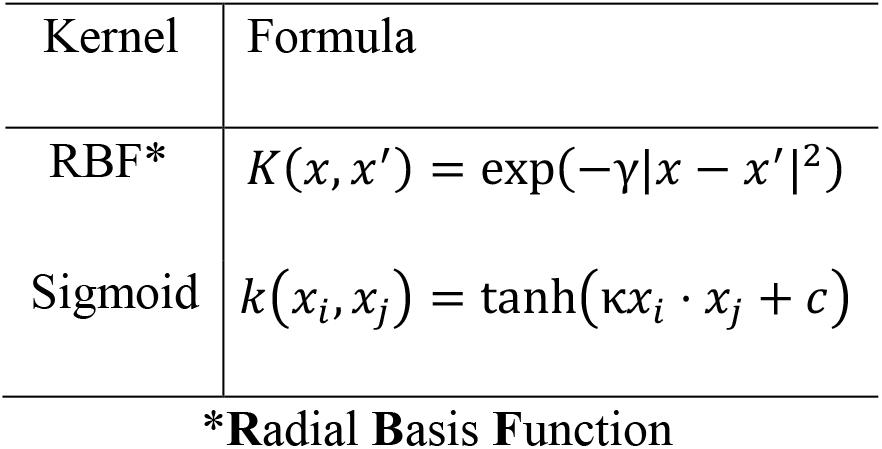
Kernel used on SVC training phase.

Finally, XGBoost [52] is a boosting technique that creates a predictive model using additive decision trees. The prediction for an instance is given by ^*yi* = ^P^*k* = 1^*K*^*f*_*k*_(*x*_*i*_), with each *f*_*k*_ being a regression tree. The model optimises an objective function by combining the empirical loss and a regularisation term to penalise the model’s complexity. The gradients and Hessians of the loss function are used to calculate *j* for each leaf. To assess the benefit from a split, the predictive quality before and after dividing a node in two is calculated accounting for gradients, Hessians, and the *γ* penalty.

Each model has been tested in a specific evaluation phase using the most commonly used state-of-the-art criteria for classification tasks. Specifically, we have calculated the accuracy, i.e. the percentage of correct predictions out of the total number of predictions defined by equation 6);

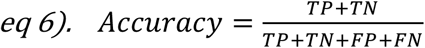

Where TP are the True Positives, TN are the True Negatives, FP are the False Positives and FN are the False Negatives.

Furthermore, we also checked the precision index that quantifies the true positive ration among all positive predictions (equation 7)

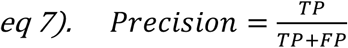

And the Recall, or Sensitivity, index which measures the ratio between the true positive predictions over the sum between true positive and false negative (equation 8), thus giving a measure of the discriminative capacity of the model

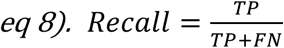

Finally, the F1-Score (equation 9), i.e. the harmonic mean of precision and recall, was considered since it provides an analytical compromise for extremely unbalanced datasets such as that under consideration

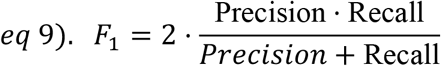

Indeed, considering its ability to evaluate highly unbalanced datasets, the F1 score was the most used metric for both feature selection and overall model performance.

As for feature selection, the number of retained features was varied from 50 to 300 in steps of 10 assessing the overall performance with F1-score.

Importantly, during the evaluation phase of the best model, accuracy was replaced by balanced accuracy (equation 10) to take into account class imbalance.

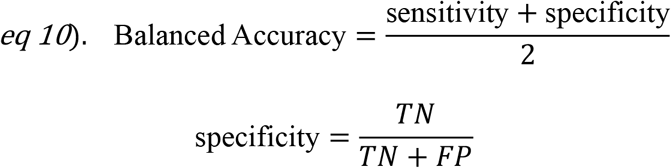

### Web Interface

The AGAPE web interface development was carried out using the server-side Django 5.2.3 (Python 3.13) framework, combined with standard front-end technologies such as HTML5, SCSS, JavaScript, and Bootstrap 5.3.6 CSS framework.

As shown in Figure 2, the interface is divided into several sections accessible via a navigation menu: a home page presenting the AGAPE project, a section dedicated to submitting molecules and their descriptors, a section for displaying results, and a section devoted to feedback and contact information. In its current version, AGAPE relies exclusively on the input of precomputed classical and quantum chemical descriptors provided by the user either through manual entry or CSV file upload.

**Fig. 2.**
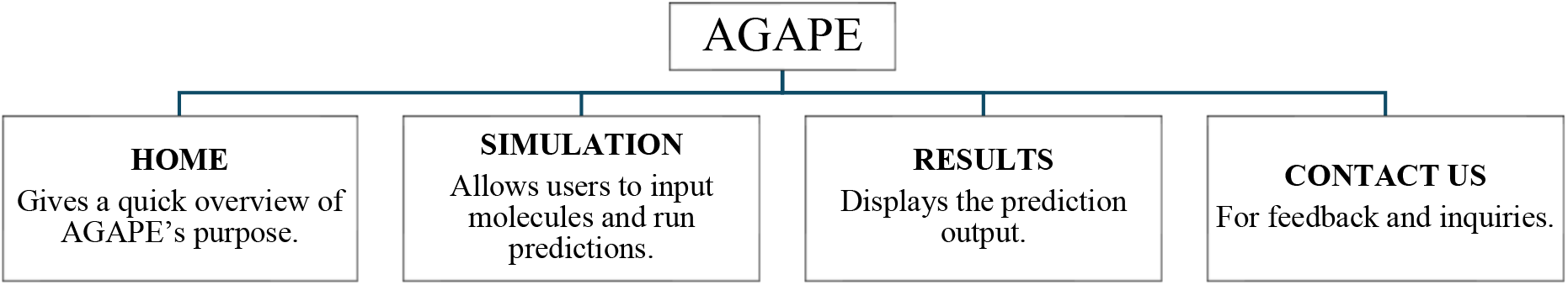
Overview of the AGAPE web interface, highlighting its different pages and their specific functions.

Supported descriptor formats include those generated by AlvaDesc (classical) and Jaguar (quantum). A downloadable example CSV file is available on the simulation page, illustrating the required column structure and formatting.

The schematic representation of the platform data flow is showed in Figure 3. Users submit input either manually or via CSV upload. Classical and quantum chemical descriptors are merged and fed into the AGAPE machine learning model, which predicts the likelihood of G4 stabilization by the given molecule.

**Fig. 3.**
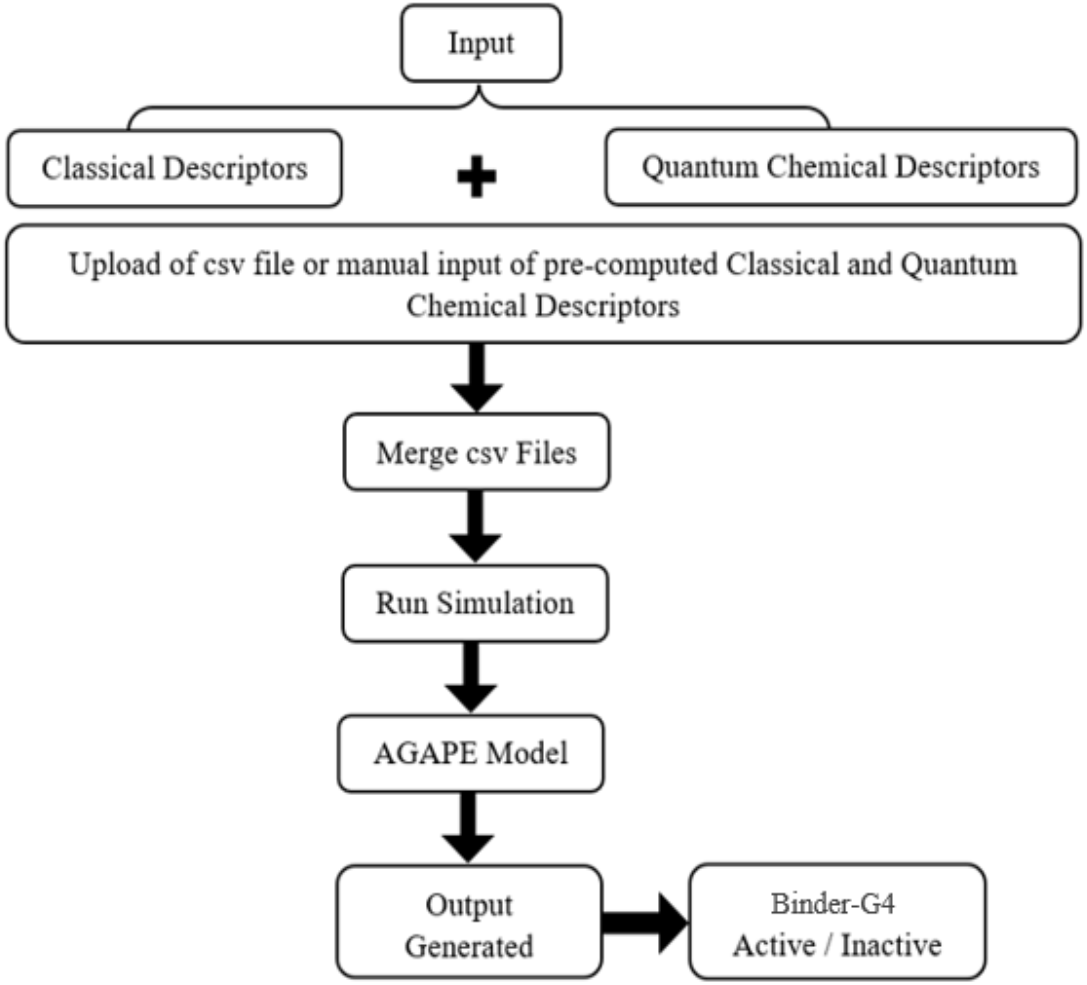
Overview of the platform’s data flow.

The results are displayed dynamically, accompanied by a classification message (active/inactive), and can also be downloaded in CSV format. The website allows batch processing, for over 1000 molecules in a single submission. Validation checkpoints have been implemented to control input formats, notably to ensure compliance with the input requirements defined by AGAPE, and to prevent submission errors. The entire system has been designed with ease of use and extensibility in mind. The platform was deployed locally using Docker 28.2.2 and Docker Compose v2.37.1, encapsulating all dependencies into isolated containers to ensure environment reproducibility and portability. For production-level testing, the application was served using Nginx 1.28, a high-performance reverse proxy and web server, providing efficient static file delivery and routing. Hosting support is provided by the Fondazione Ri.MED server infrastructure, enabling secure access for collaborative development and remote evaluation.

From a data privacy standpoint, no user data is stored permanently on the server. All files uploaded by users are temporarily processed during the session, strictly for the purpose of model inference, and are automatically deleted immediately after analysis. This ensures full compliance with data confidentiality best practices and minimizes the risk of residual data exposure.

An upcoming version of the AGAPE platform aims to automate the generation of classical descriptors using an integrated Python-based cheminformatics library. This functionality will be triggered either by direct submission of a SMILES string or via a molecule drawn using the JSME molecular editor, an interactive JavaScript-based tool embedded within the browser. The editor will automatically translate the drawn structure into its corresponding SMILES representation, which is then used to compute descriptors and feed the model. However, this automatization will require avoiding the use of commercial software for the prediction of the descriptors and will thus require a retraining of the model.

## Results and Discussion

This section will describe the results obtained during the different stages of our work. Specifically, the results obtained with the various Feature Selection methods will be described, followed by the results obtained from training the best model for the selected feature subsets.

### Dataset creation

The latest version of G4LDB includes 3695 G4 binders and 32142 activity entries. From this database, we focused on interaction and stabilization activities, specifically selecting data related to the FRET melting assay, since the latter is a widely used technique to measure the melting temperature (T_m_) of G4 structures, providing insights into their thermal stability and the stabilization induced by small molecules.

In a typical FRET assay, a G4 sequence is labeled at its 5’ and 3’ ends with fluorescent donor and acceptor dyes. Upon excitation of the donor, energy transfer occurs if the acceptor is in close spatial proximity, thus quenching the donor fluorescence. FRET efficiency, because of the high sensitivity to the acceptor and donor distance R because of its R^-6^ dependence, is an effective tool for assessing G4 folding and unfolding. At lower temperatures, G4s remain folded, keeping the donor and acceptor dyes close enough for FRET to occur. As the temperature rises, the G4 unfolds, increasing the distance between the dyes, reducing FRET efficiency, and enhancing the donor fluorescence quantum yield. In the presence of a stabilizing compound, the FRET assay is repeated. A stabilizer increases the T_m_ of the G4, indicating that the structure remains intact at higher temperatures. The shift in melting temperature (ΔT_m_) provides a direct measure of the stabilization induced by the binder. We selected this technique for its consistency with our in-house dataset, which includes stabilization data obtained from FRET assays on various G4-DNA sequences. After filtering duplicates from G4LDB, we obtained 5320 unique activity entries and 1835 unique binders. Additional filtering ensured consistency of T_m_ data for molecules with multiple activity records. To align with our research group’s focus on metal complexes, particularly salphen-based ligands known for their G4 stabilization activity [10, 12], we supplemented the G4LDB collection with annotated salphen-like complexes from literature sources [53–56].

Ultimately, we compiled a unique dataset comprising 1217 compounds, categorized into 490 ACTIVE and 727 INACTIVE entries. Among these, 1073 were purely organic compounds, while 144 were metal complexes. Following QC geometry optimization and property calculations, the final dataset included 1188 molecules.

### Descriptors calculation

For each molecule, a total of 5666 molecular descriptors spanning 33 distinct classes were calculated using alvaDesc.[48] These classes include Topological Indices, Connectivity Indices, Geometrical Descriptors, 3D Autocorrelations, Functional Group Counts, Charge Descriptors, Molecular Properties, Drug-like Indices, and others. Detailed information about the descriptors can be found at https://www.alvascience.com/alvadesc-descriptors.

A preliminary dimensionality reduction step was performed to streamline the dataset. Features with constant values, standard deviations below 0.0001, or columns containing all missing values were removed. This filtering process reduced the dataset to 4285 conserved features from alvaDesc.

Additionally, relevant quantum chemical (QC) properties were computed through Density Functional Theory (DFT) calculations. These included descriptors such as: i) Surfaces: electrostatic potential, average local ionization energy, and electron densities; ii) Atomic Electrostatic Potential Charges: Charges and dipole moments; iii) Electronic Properties: Mulliken population, multipole moments, polarizability, and hyperpolarizability. The QC descriptors provide critical insights into molecular and electronic properties, complementing the alvaDesc features to enhance the predictive capabilities of the model.

### Feature Selection

#### Filter and Embedded FS

This section shows a summary table that collects the main results involving the selection of the most preminent features. Note that additional data are also presented in the supplementary material (Table S1-S11). Table 2 reports the top-performing model identified for each feature selection strategy used in the study using an 80:10:10 split train-validation-test.

**Table 2.** Summary table on test set split 80:10:10.

| Model | SS* | Accuracy | Precision | Recall | F1 | # features |
| --- | --- | --- | --- | --- | --- | --- |
| DT | Mutual Info | 0.8067 | 0.7551 | 0.7708 | 0.7629 | 140 |
| DT | ANOVA | 0.7983 | 0.7608 | 0.7291 | 0.7446 | 150 |
| DT | Chi2 | 0.7899 | 0.7555 | 0.7083 | 0.7311 | 180 |
| NB | Mutual Info | 0.6891 | 0.6000 | 0.6875 | 0.6408 | 100 |
| NB | ANOVA | 0.6387 | 0.5490 | 0.5833 | 0.5657 | 190 |
| NB | Chi2 | 0.6891 | 0.5846 | 0.7917 | 0.6726 | 210 |
| RF | Mutual Info | 0.8319 | 0.8333 | 0.7292 | 0.7778 | 260 |
| RF | ANOVA | 0.8151 | 0.7955 | 0.7292 | 0.7609 | 60 |
| RF | Chi2 | 0.8319 | 0.8500 | 0.7083 | 0.7727 | 70 |
| RF | RFI** | 0.8151 | 0.8250 | 0.6875 | 0.7500 | 260 |

\*SS = Selection Strategy; \*\*RFI = RandomForestImportance

From this data it appears that the most efficient model is RF combined with Mutual Information as future selection method, as it achieved the highest F1 score and, consequently, the highest accuracy. Table 3 summarises the combinations of feature selection methods in cross-validation.

**Table 3.** Summary table on cross validation.

| Model | SS* | Accuracy | Precision | Recall | F1 | # features |
| --- | --- | --- | --- | --- | --- | --- |
| DT | Mutual Info | 0.7972 | 0.7529 | 0.7428 | 0.7453 | 120 |
| DT | ANOVA | 0.7937 | 0.7530 | 0.7238 | 0.7352 | 240 |
| DT | Chi2 | 0.7862 | 0.7217 | 0.7564 | 0.7368 | 130 |
| NB | Mutual Info | 0.6826 | 0.5984 | 0.6549 | 0.6196 | 160 |
| NB | ANOVA | 0.6919 | 0.6112 | 0.6461 | 0.6231 | 150 |
| NB | Chi2 | 0.6170 | 0.5200 | 0.8573 | 0.6421 | 290 |
| RF | MutualInfo | 0.8636 | 0.8432 | 0.8081 | 0.8237 | 300 |
| RF | ANOVA | 0.8586 | 0.8473 | 0.7908 | 0.8150 | 290 |
| RF | Chi2 | 0.8611 | 0.8400 | 0.8044 | 0.8204 | 60 |
| RF | RFI** | 0.8636 | 0.8470 | 0.8054 | 0.8236 | 100 |

\*SS = Selection Strategy; \*\*RFI = RandomForestImportance

The results highlight that RF consistently achieved better performance reaching out F1 = 0.8236 when using the embedded method.

#### Wrapper FS

As described in the methods section, the selection of features with wrapper methods was conducted using Sequential Forward Selection with a maximum number of features of 1723. Indeed, to perform an exhaustive search of the features, it was essential to search within the entire d-dimensional space. Moreover, to ensure that all possible data distributions are tested, the cross validation parameter (*cv* = 10) has been set. In this way, feature selection was done using the average across folds to get a robust result that was invariant to different distributions. Afterwards, once the parameters of the SFS algorithm set, all the selected ML models have been tested. The results are shown in Figure 4.

**Fig. 4.**
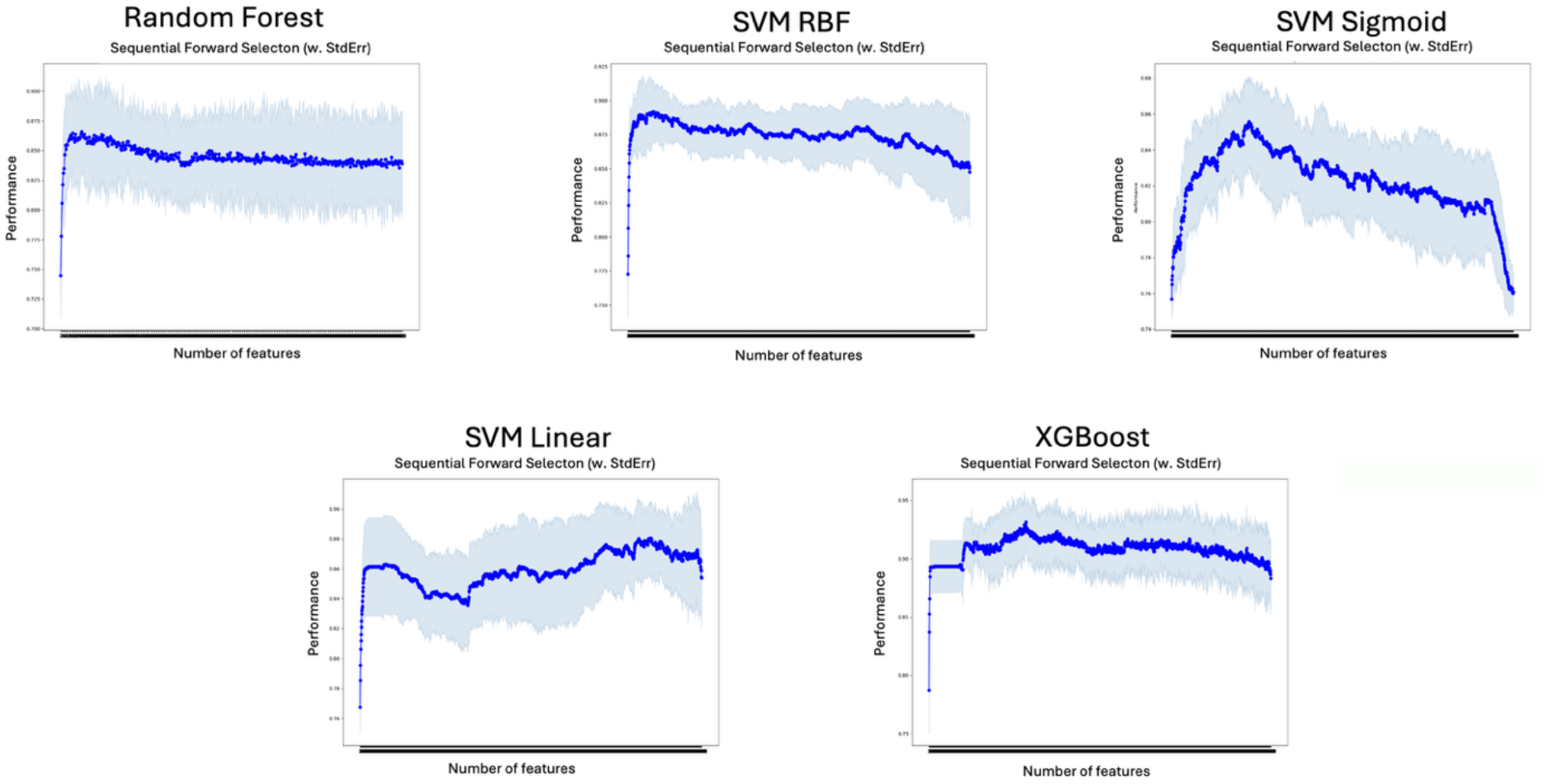
Plot showing comparative results obtained with various ML approaches in Sequential Feature Selection (SFS) on the total feature set.

The overall trend observed in the various feature selection studies is consistent with what has been reported in the literature. Indeed, the increase in performance is not directly related to the number of features utilised. However, as shown in Figure 4, it decreases for almost all models as the number of features increases. The best stability was obtained with the XGBoost model, which follows the trend discussed above but achieves higher performance in terms of F1-score compared to the other models. As can be seen from Figure 5, a peak performance of ∼ 0.93 is achieved.

**Fig. 5.**
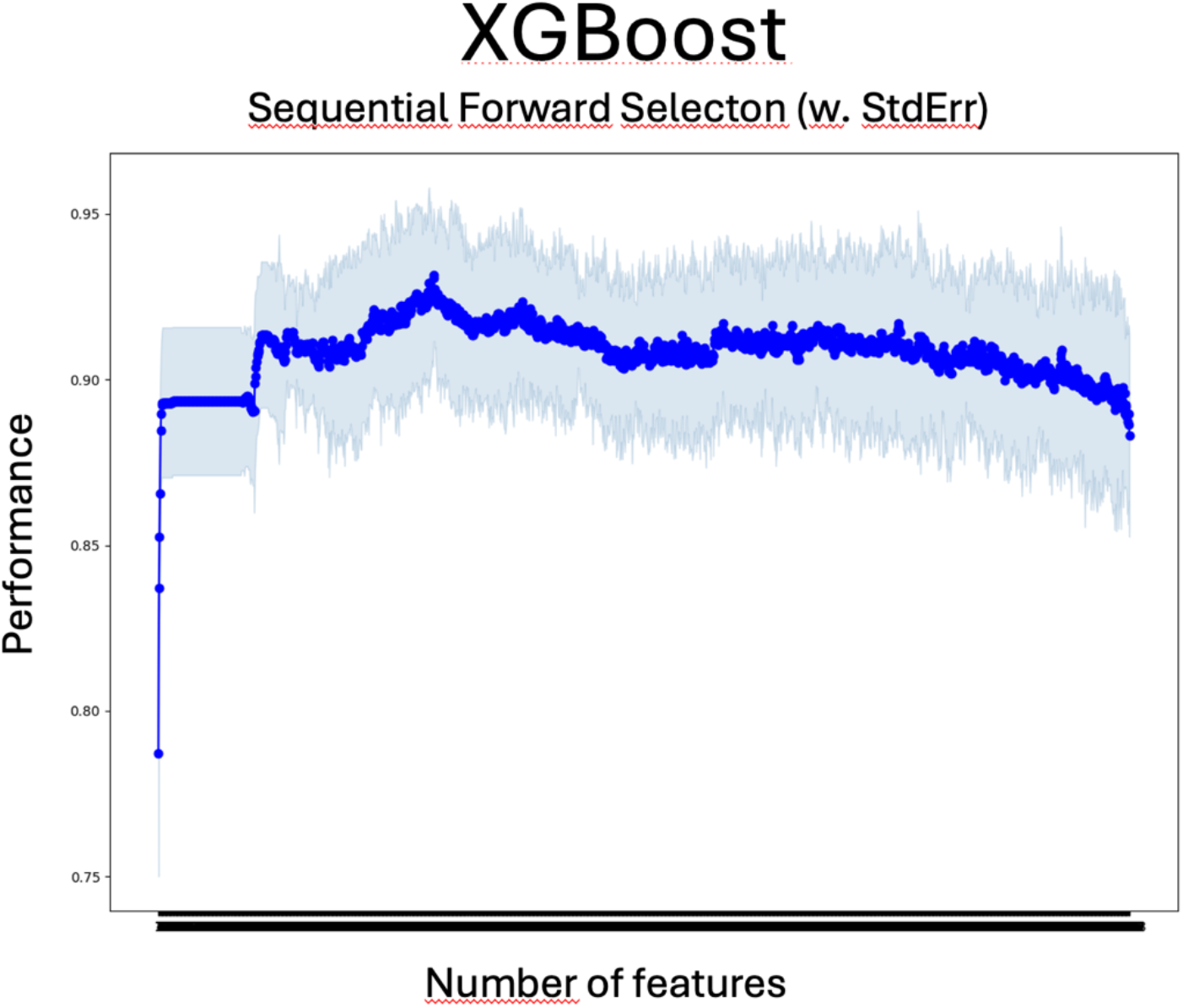
Plot of the SFS procedure on XGBoost.

XGBoost outperforms the models’ results regardless of the type of feature selection used. The highest performance was achieved using a collection of 489 features which reduced the dataset’s dimensionality by one-third. To further minimise data complexity, an additional subset of 180 features (one-tenth of the total) was selected still yielding acceptable F1-scores. These two features set were then used to carry out the classification tests described in the following section.

### Classification

The classification trials had been carried out using XGBoost in its best configuration, using the two subsets of features identified in the previous stages. To maximise the performance of XGBoost, hyperparameter tuning was performed by evaluating several parameters combinations. The optimum configuration was determined and is reported in Table 4.

**Table 4.** Summary of the best parameters used for training XGBoost.

| Parameter | Values | Description |
| --- | --- | --- |
| n estimators | 1000 | Total number of trees (boosting rounds).<br>A high value allows for more gradual learning, which is useful with a low eta |
| max depth | 8 | Maximum depth of trees. Higher values allow for more complex models, but increase the risk of overfitting |
| eval metric | auc | Evaluation metric: AUC (Area Under the Curve), useful for evaluating the discriminatory capacity of the model |
| booster | dart | Type of booster used. dart (Dropouts meet Multiple Additive Regression Trees), is a boosting method that adds dropout to trees during training to reduce overfitting and improve generalization. |
| eta | 0.05 | Learning rate |
| subsample | 1 | Percentage of samples used for each tree. 1 means that all data are used (no sub-sampling) |
| scale_pos_weight | 3 | Weight assigned to the positive class. Useful for managing unbalanced classes |

The trials used 10-fold cross-validation to assess the algorithm’s robustness. Tables 5 and 6 show the results obtained for the two feature subsets (489 and 180) for all 10 folds; both tables indicate the average performance over the 10 folds evaluated on the last row.

**Table 5.** Results obtained with XGBoost on larger (489) subset of feature.

| FOLD | F1 | BAL. ACC. | PRECISION | RECALL |
| --- | --- | --- | --- | --- |
| 1 | 0.9116 | 0.8802 | 0.8933 | 0.9306 |
| 2 | 0.9241 | 0.9014 | 0.9178 | 0.9306 |
| 3 | 0.9091 | 0.8848 | 0.9028 | 0.9155 |
| 4 | 0.9091 | 0.8575 | 0.8434 | 0.9859 |
| 5 | 0.9020 | 0.8505 | 0.8415 | 0.9718 |
| 6 | 0.8767 | 0.8361 | 0.8533 | 0.9014 |
| 7 | 0.9577 | 0.9476 | 0.9577 | 0.9577 |
| 8 | 0.9116 | 0.8781 | 0.8816 | 0.9437 |
| 9 | 0.8874 | 0.8335 | 0.8375 | 0.9437 |
| 10 | 0.8844 | 0.8407 | 0.8553 | 0.9155 |
|  | *Avg F1* | *Avg Bal. Acc.* | *Avg Precision* | *Avg Recall* |
|  | **0.9074** | **0.8711** | **0.8784** | **0.9396** |

**Table 6.** Results obtained with XGBoost on smaller (180) subset of feature.

| FOLD | F1 | BAL. ACC. | PRECISION | RECALL |
| --- | --- | --- | --- | --- |
| 1 | 0.8811 | 0.8524 | 0.8873 | 0.8750 |
| 2 | 0.9333 | 0.9010 | 0.8974 | 0.9722 |
| 3 | 0.9209 | 0.9090 | 0.9412 | 0.9014 |
| 4 | 0.8961 | 0.8401 | 0.8313 | 0.9718 |
| 5 | 0.8774 | 0.8122 | 0.8095 | 0.9577 |
| 6 | 0.8611 | 0.8220 | 0.8493 | 0.8732 |
| 7 | 0.9379 | 0.9164 | 0.9189 | 0.9577 |
| 8 | 0.8811 | 0.8499 | 0.8750 | 0.8873 |
| 9 | 0.8552 | 0.8090 | 0.8378 | 0.8732 |
| 10 | 0.8732 | 0.8409 | 0.8732 | 0.8732 |
|  | *Avg F1* | *Avg Bal. Acc.* | *Avg Precision* | *Avg Recall* |
|  | **0.8917** | **0.8553** | **0.8721** | **0.9143** |

As shown from the average performances reported in Tables 5 and 6, both subsets consistently yielded good performances for each of the 10 folds calculated. The use of a higher number of features led to a peak sensitivity of 0.9396, proving the model’s ability to correctly discriminate truly active molecules. The performance is encouraging, especially considering the small number of training samples, even though they were tested exhaustively using 10-fold cross-validation.

In order to assess deeper the accuracy of the model, an additional test data set consisting of 28 molecules was selected and used to validate the best algorithm. The results obtained from both subsets are shown in Table 7. On the test set, the performance of our model drops down. The cause behind this lack of performance could be ascribed to different reasons, not necessarily connected to the model effectiveness, but rather due to statistical and sampling limitations inherent in small datasets used as external set. In this case, our external set mainly contains inorganic compounds that were less represented into the initial dataset, thus leading to a biased estimate of the model generalisation. Indeed, the model may analyse unfamiliar patterns or chemotypes probably without having enough context to generalize, leading to degraded performance.

**Table 7.** Results obtained on the Test data.

| # FEATURES | F1 | BAL. ACC. | PRECISION | RECALL | AUC |
| --- | --- | --- | --- | --- | --- |
| **489** | 0.7357 | 0.6661 | 0.7832 | 0.7222 | 0.6661 |
| **180** | 0.6928 | 0.6644 | 0.6972 | 0.6889 | 0.6644 |

#### SHAP analysis

SHAP [57] is an Explainable AI approach based on game theory to explain the output and mechanisms underlying the prediction of a Machine Learning model. Specifically, SHAP computes the Shapley values for a given instance according to the equation 8):

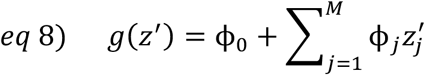

where *g* is the explanation of the model, *M* is the maximum simplified input features size contained in *z*^′^, and *ϕ*_*i*_ ∈ R.

In our case, we used a variant of the SHAP explainer, namely TreeExplainer [58], which is designed to interpret tree-based ML models such as Decision Tree, Random Forest, and Gradient-Boosted Trees.

The results obtained from this analysis are shown in Figure 6, which displays a ViolinPlot of the top 10 most relevant features.

**Fig. 6.**
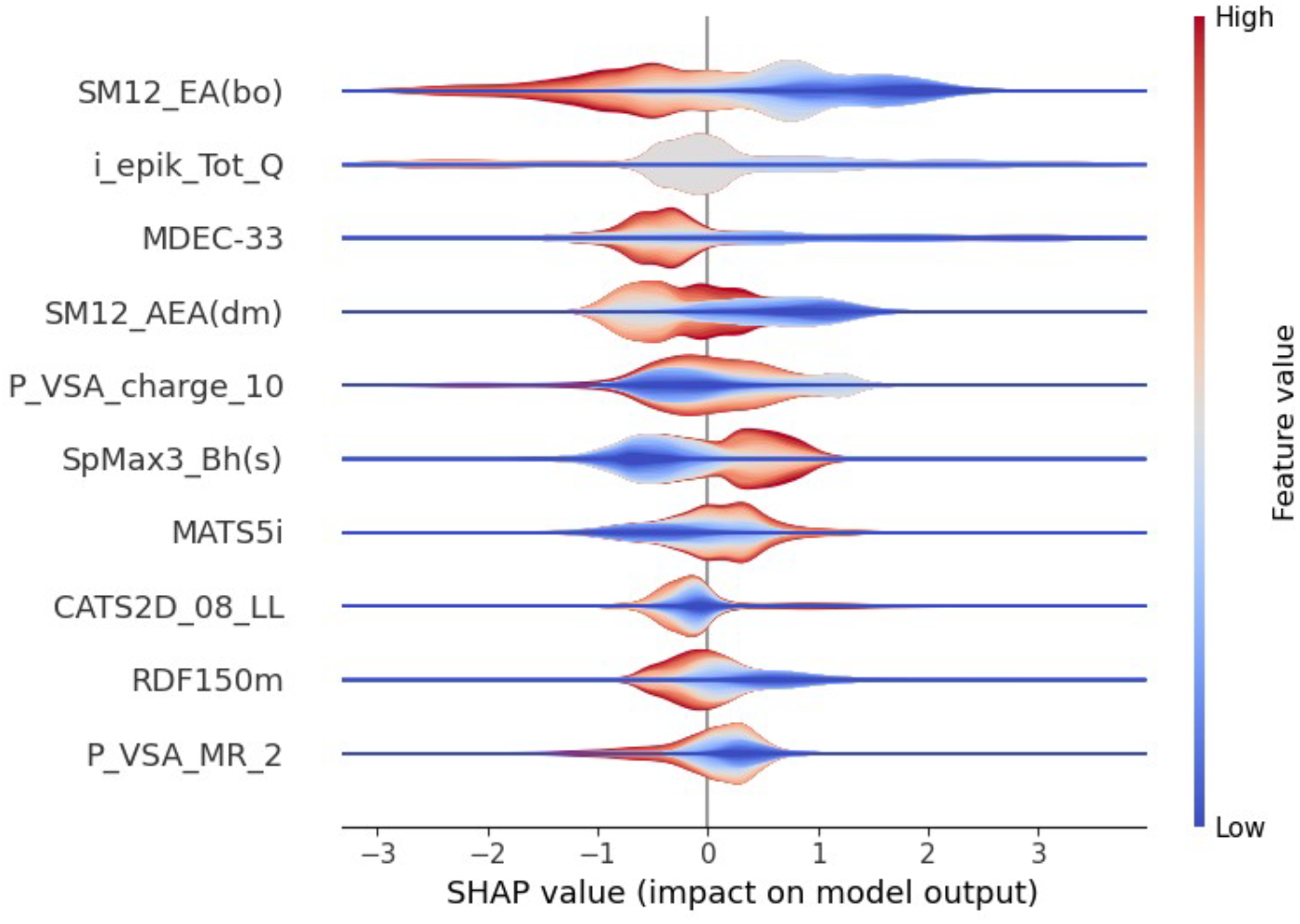
Violinplot that reports the contribution of the different feature obtained with the SHAP analysis This kind of descriptors are particularly used for QSAR application and property prediction of small molecules because they consider the whole molecular connectivity and atomic feature at the same time.

The relevance of each individual feature can be interpreted by evaluating the value of the individual Shapley values obtained. Specifically, values less than 0 indicate a negative impact on the prediction, while values greater than 0 suggest a positive contribution. Based on this assumption, we can see that the feature SM12 EA (bo) (spectral moment of order 12 from edge adjacency mat. weighted by bond order) tends to negatively influence the correct classification when it assumes high values. This descriptor considers long-range interactions within molecular topology and it could some potential molecular rigidity and extensive conjugation. A very high value of this descriptor could refer to molecular complexity and rigidity which might hinder molecule ability to adopt an effective conformation to bind and stabilise G4 conformation through a π–π stacking or groove binding in G4 DNA. Molecules with excessively complex or branched topologies could indeed present steric hindrance or lack the surface planarity required for optimal binding.

In contrast, SM12 AEA(dm) (spectral moment of order 12 from augmented edge adjacency mat. weighted by dipole moment), SpMax3 Bh(s) (largest eigenvalue n. 3 of Burden matrix weighted by I-state), are among those that positively influence classification performance when their values are high. SM12 AEA(dm) is a variation of the previous descriptor, but considering polarity distribution into the topological structure. A favourable polar distribution across the molecular surface is crucial to stabilise the G4 conformation and the polarity of the molecule could facilitate the binding process through polar interactions (i.e. hydrogen-bonding interactions). SpMax3 Bh(s) reflects electronic distribution and atomic hybridization within the molecule. High values of this descriptor could be related with delocalized π-electrons (aromatic systems or π conjugation), a favourable molecular characteristic to stabilize DNA G4 conformation through π–π interactions [29].

Of particular interest, then, is the behaviour of the i epik Tot Q (Net charge of the molecule) feature, which turns out to be almost irrelevant in determining the classification features. This result is comforting because its value is set to 0 for all samples, given the complex computation of this feature. This choice was made to retain the feature in the optimal feature set, ensuring the possibility of reintroducing it in future model re-training stages. Net molecular charge is not a significant parameter to explain binding affinity to G4. It is well known, indeed, that neutral ligands are effective in G4 binding and stabilisation process.

The negligible impact of this feature provides further evidence of the robustness of the approach in accurately assessing the contribution of each variable. As shown by our results, the most influential features in distinguishing active from inactive molecules are those related to molecular topology, molecular polarity and π conjugation, particularly the edge adjacency indices and Burden eigenvalues. Both descriptors effectively capture the global structure of the molecule while incorporating information about its electronic properties [52, 59, 60].

## Conclusions

In this work, we introduced AGAPE (computational G-quadruplex Affinity Prediction), the first *in silico* platform designed to predict the affinity and stabilization capacity of G4 binders. AGAPE employs a machine learning framework based on supervised classification, using molecular descriptors and quantum chemical derived properties of small molecules. To develop and train the models, we curated a dataset of 476 ACTIVE and 712 INACTIVE compounds, enabling the identification of chemical features critical to G4 stabilization.

Among the tested classifiers, XGBoost demonstrated the best predictive performance, achieving an average F1-score of 0.91 and a peak sensitivity of 0.94 in cross-validation, confirming its strong generalization ability even with a limited number of training samples. The robustness of the model was further validated on an independent in-house library of 28 compounds, which included 25 transition metal complexes and 3 organic ligands not included in the training, test, or validation sets. Despite the inorganic compounds representing the minority class in the original dataset, the model achieved an overall accuracy of 66%. Notably, feature importance analysis using SHAP provided interpretable insights, confirming the relevance of molecular topology and electronic structure descriptors—particularly edge adjacency indices and Burden eigenvalues—in determining G4 binding potential.

The AGAPE framework is deployed through an accessible and secure web interface, allowing researchers to perform batch predictions based on user-supplied molecular descriptors. This platform not only enables high-throughput screening of potential G4 binders but also lays the foundation for integrating cheminformatics and quantum-informed descriptors in predictive modelling.

Overall, AGAPE offers a novel, interpretable, and practical tool for accelerating the discovery of selective G4-targeting compounds. Future work will focus on expanding the dataset, automating descriptor calculation from SMILES input. We aim to refine AGAPE expanding the model to predict selectivity for G4 structures over canonical nucleic acids, as well as selectivity across different G4 topologies or sequences, represents an exciting direction for further development. Additionally, integrating AGAPE with generative AI frameworks could enable the suggestion of structural modifications to existing G4 stabilizers or the design of novel scaffolds. By virtually labeling large datasets as ACTIVE or INACTIVE for G4 stabilization, AGAPE could provide the foundation for training generative AI or large language models (LLMs). These tools could mitigate the scarcity of experimental data and drive the discovery of new, selective G4-targeting compounds.

## Declaration

### Author contributions

Luisa D’Anna: conceptualization, dataset creation, formal analysis, visualization, writing – original draft; Salvatore Contino: ML methodology, formal analysis, visualization, writing – original draft; Rosalinda Marinello: ML methodology, formal analysis, visualization, writing – original draft; Julie Fares: Webapp creation, formal analysis, visualization, writing – original draft; Giada De Simone: dataset creation; Florent Barbault: Webapp creation, formal analysis, visualization, writing – original draft; Antonio Monari: conceptualization, writing – original draft, formal analysis, visualization; Giampaolo Barone: conceptualization, writing – original draft, formal analysis, visualization; Alessio Terenzi: conceptualization, writing – original draft, formal analysis, visualization. Ugo Perricone: conceptualization, writing – original draft, formal analysis, visualization.

### Data availability

#### WebApp addres

http://agape.fondazionerimed.com:4444.

Dataset and ML models are available at https://github.com/Molinf-RiMED/AGAPE https://doi.org/10.5281/zenodo.17277995

