## Supplemental tables for "AGAPE (computAtional G-quadruplex Affinitiy PrEdiction): The first Artificial Intelligence workflow for G-quadruplex binding affinity prediction"

**Supplementary material**

In this section it show a different result for different selection of features. The best model are in bold.

### 1.1 SPLIT 80:10:10

This subsection presents results using a train-test split with 80% training, 10% validation and 10% testing.

| Model | SS | Accuracy | Precision | Recall | F1 | # features |
|---|---|---|---|---|---|---|
| RF | Mutual Info | 0.7983 | 0.7727 | 0.7083 | 0.7391 | 50 |
| RF | Mutual Info | 0.7983 | 0.8000 | 0.6667 | 0.7273 | 60 |
| RF | Mutual Info | 0.8151 | 0.8095 | 0.7083 | 0.7556 | 70 |
| RF | Mutual Info | 0.8151 | 0.8095 | 0.7083 | 0.7556 | 80 |
| RF | Mutual Info | 0.8487 | 0.8571 | 0.7500 | 0.8000 | 90 |
| RF | Mutual Info | 0.8067 | 0.8049 | 0.6875 | 0.7416 | 100 |
| RF | Mutual Info | 0.7983 | 0.7857 | 0.6875 | 0.7333 | 110 |
| RF | Mutual Info | 0.8235 | 0.8293 | 0.7083 | 0.7640 | 120 |
| RF | Mutual Info | 0.8235 | 0.8293 | 0.7083 | 0.7640 | 130 |
| RF | Mutual Info | 0.8235 | 0.8140 | 0.7292 | 0.7692 | 140 |
| RF | Mutual Info | 0.7899 | 0.7949 | 0.6458 | 0.7126 | 150 |
| RF | Mutual Info | 0.7899 | 0.7805 | 0.6667 | 0.7191 | 160 |
| RF | Mutual Info | 0.7983 | 0.7857 | 0.6875 | 0.7333 | 170 |
| RF | Mutual Info | 0.8151 | 0.8250 | 0.6875 | 0.7500 | 180 |
| RF | Mutual Info | 0.7899 | 0.7805 | 0.6667 | 0.7191 | 190 |
| RF | Mutual Info | 0.7983 | 0.7857 | 0.6875 | 0.7333 | 200 |
| RF | Mutual Info | 0.7899 | 0.7805 | 0.6667 | 0.7191 | 210 |
| RF | Mutual Info | 0.7647 | 0.7500 | 0.6250 | 0.6818 | 220 |
| RF | Mutual Info | 0.8067 | 0.8049 | 0.6875 | 0.7416 | 230 |
| RF | Mutual Info | 0.8067 | 0.8205 | 0.6667 | 0.7356 | 240 |
| RF | Mutual Info | 0.8235 | 0.8293 | 0.7083 | 0.7640 | 250 |
| **RF** | **Mutual Info** | **0.8319** | **0.8333** | **0.7292** | **0.7778** | **260** |
| RF | Mutual Info | 0.8067 | 0.8205 | 0.6667 | 0.7356 | 270 |
| RF | Mutual Info | 0.8151 | 0.8250 | 0.6875 | 0.7500 | 280 |
| RF | Mutual Info | 0.8067 | 0.7907 | 0.7083 | 0.7473 | 290 |
| RF | Mutual Info | 0.8067 | 0.8049 | 0.6875 | 0.7416 | 300 |

Table 1: Performance of the Random Forest model with Mutual Information selection on different numbers of features.

| Model | SS | Accuracy | Precision | Recall | F1 | # features |
|---|---|---|---|---|---|---|
| RF | ANOVA | 0.7815 | 0.7895 | 0.6250 | 0.6977 | 50 |
| **RF** | **ANOVA** | **0.8151** | **0.7955** | **0.7292** | **0.7609** | **60** |
| RF | ANOVA | 0.7983 | 0.8000 | 0.6667 | 0.7273 | 70 |
| RF | ANOVA | 0.7983 | 0.7857 | 0.6875 | 0.7333 | 80 |
| RF | ANOVA | 0.7983 | 0.7857 | 0.6875 | 0.7333 | 90 |
| RF | ANOVA | 0.7731 | 0.7442 | 0.6667 | 0.7033 | 100 |
| RF | ANOVA | 0.8067 | 0.7907 | 0.7083 | 0.7473 | 110 |
| RF | ANOVA | 0.7647 | 0.7381 | 0.6458 | 0.6889 | 120 |
| RF | ANOVA | 0.7899 | 0.7674 | 0.6875 | 0.7253 | 130 |
| RF | ANOVA | 0.7815 | 0.7750 | 0.6458 | 0.7045 | 140 |
| RF | ANOVA | 0.7815 | 0.7619 | 0.6667 | 0.7111 | 150 |
| RF | ANOVA | 0.7731 | 0.7442 | 0.6667 | 0.7033 | 160 |
| RF | ANOVA | 0.7647 | 0.7632 | 0.6042 | 0.6744 | 170 |
| RF | ANOVA | 0.7647 | 0.7500 | 0.6250 | 0.6818 | 180 |
| RF | ANOVA | 0.7647 | 0.7381 | 0.6458 | 0.6889 | 190 |
| RF | ANOVA | 0.7899 | 0.7949 | 0.6458 | 0.7126 | 200 |
| RF | ANOVA | 0.7899 | 0.7674 | 0.6875 | 0.7253 | 210 |
| RF | ANOVA | 0.7647 | 0.7500 | 0.6250 | 0.6818 | 220 |
| RF | ANOVA | 0.7647 | 0.7500 | 0.6250 | 0.6818 | 230 |
| RF | ANOVA | 0.7647 | 0.7632 | 0.6042 | 0.6744 | 240 |
| RF | ANOVA | 0.7731 | 0.7561 | 0.6458 | 0.6966 | 250 |
| RF | ANOVA | 0.7563 | 0.7436 | 0.6042 | 0.6667 | 260 |
| RF | ANOVA | 0.7815 | 0.7895 | 0.6250 | 0.6977 | 270 |
| RF | ANOVA | 0.7815 | 0.7619 | 0.6667 | 0.7111 | 280 |
| RF | ANOVA | 0.7647 | 0.7381 | 0.6458 | 0.6889 | 290 |
| RF | ANOVA | 0.7647 | 0.7632 | 0.6042 | 0.6744 | 300 |

Table 2: Performance of the Random Forest model with ANOVA seldifferent numbers of features.

| Model | SS | Accuracy | Precision | Recall | F1 | # features |
|---|---|---|---|---|---|---|
| RF | Chi2 | 0.7815 | 0.8056 | 0.6042 | 0.6905 | 50 |
| RF | Chi2 | 0.7983 | 0.8529 | 0.6042 | 0.7073 | 60 |
| **RF** | **Chi2** | **0.8319** | **0.8500** | **0.7083** | **0.7727** | **70** |
| RF | Chi2 | 0.8235 | 0.8293 | 0.7083 | 0.7640 | 80 |
| RF | Chi2 | 0.7983 | 0.7857 | 0.6875 | 0.7333 | 90 |
| RF | Chi2 | 0.8067 | 0.8378 | 0.6458 | 0.7294 | 100 |
| RF | Chi2 | 0.7731 | 0.7692 | 0.6250 | 0.6897 | 110 |
| RF | Chi2 | 0.8067 | 0.8049 | 0.6875 | 0.7416 | 120 |
| RF | Chi2 | 0.7983 | 0.8000 | 0.6667 | 0.7273 | 130 |
| RF | Chi2 | 0.7983 | 0.7857 | 0.6875 | 0.7333 | 140 |
| RF | Chi2 | 0.7815 | 0.8056 | 0.6042 | 0.6905 | 150 |
| RF | Chi2 | 0.7983 | 0.7857 | 0.6875 | 0.7333 | 160 |
| RF | Chi2 | 0.8067 | 0.7907 | 0.7083 | 0.7473 | 170 |
| RF | Chi2 | 0.7899 | 0.7805 | 0.6667 | 0.7191 | 180 |
| RF | Chi2 | 0.8067 | 0.8049 | 0.6875 | 0.7416 | 190 |
| RF | Chi2 | 0.8067 | 0.8205 | 0.6667 | 0.7356 | 200 |
| RF | Chi2 | 0.7815 | 0.7619 | 0.6667 | 0.7111 | 210 |
| RF | Chi2 | 0.8067 | 0.8205 | 0.6667 | 0.7356 | 220 |
| RF | Chi2 | 0.7983 | 0.7857 | 0.6875 | 0.7333 | 230 |
| RF | Chi2 | 0.7983 | 0.8000 | 0.6667 | 0.7273 | 240 |
| RF | Chi2 | 0.7899 | 0.7805 | 0.6667 | 0.7191 | 250 |
| RF | Chi2 | 0.7983 | 0.8000 | 0.6667 | 0.7273 | 260 |
| RF | Chi2 | 0.8067 | 0.8205 | 0.6667 | 0.7356 | 270 |
| RF | Chi2 | 0.7983 | 0.7857 | 0.6875 | 0.7333 | 280 |
| RF | Chi2 | 0.7983 | 0.8000 | 0.6667 | 0.7273 | 290 |
| RF | Chi2 | 0.8067 | 0.8049 | 0.6875 | 0.7416 | 300 |

Table 3: Performance of the Random Forest model with Chi2 selection on different numbers of features.

| Model | SS | Accuracy | Precision | Recall | F1 | # features |
|---|---|---|---|---|---|---|
| RF | RFI | 0.7815 | 0.7500 | 0.6875 | 0.7174 | 60 |
| RF | RFI | 0.7647 | 0.7381 | 0.6458 | 0.6889 | 70 |
| RF | RFI | 0.7983 | 0.7857 | 0.6875 | 0.7333 | 80 |
| RF | RFI | 0.7983 | 0.8000 | 0.6667 | 0.7273 | 90 |
| RF | RFI | 0.7983 | 0.7727 | 0.7083 | 0.7391 | 100 |
| RF | RFI | 0.8067 | 0.8049 | 0.6875 | 0.7416 | 110 |
| RF | RFI | 0.8067 | 0.7907 | 0.7083 | 0.7473 | 120 |
| RF | RFI | 0.7899 | 0.7805 | 0.6667 | 0.7191 | 130 |
| RF | RFI | 0.7899 | 0.7674 | 0.6875 | 0.7253 | 140 |
| RF | RFI | 0.7899 | 0.7805 | 0.6667 | 0.7191 | 150 |
| RF | RFI | 0.7899 | 0.7674 | 0.6875 | 0.7253 | 160 |
| RF | RFI | 0.8067 | 0.7907 | 0.7083 | 0.7473 | 170 |
| RF | RFI | 0.7899 | 0.7674 | 0.6875 | 0.7253 | 180 |
| RF | RFI | 0.7647 | 0.7381 | 0.6458 | 0.6889 | 190 |
| RF | RFI | 0.8067 | 0.8049 | 0.6875 | 0.7416 | 200 |
| RF | RFI | 0.7899 | 0.7805 | 0.6667 | 0.7191 | 210 |
| RF | RFI | 0.7647 | 0.7500 | 0.6250 | 0.6818 | 220 |
| RF | RFI | 0.7983 | 0.7727 | 0.7083 | 0.7391 | 230 |
| RF | RFI | 0.7815 | 0.7750 | 0.6458 | 0.7045 | 240 |
| RF | RFI | 0.8067 | 0.7907 | 0.7083 | 0.7473 | 250 |
| **RF** | **RFI** | **0.8151** | **0.8250** | **0.6875** | **0.7500** | **260** |
| RF | RFI | 0.7983 | 0.7857 | 0.6875 | 0.7333 | 270 |
| RF | RFI | 0.8067 | 0.8049 | 0.6875 | 0.7416 | 280 |
| RF | RFI | 0.7815 | 0.7895 | 0.6250 | 0.6977 | 290 |
| RF | RFI | 0.8067 | 0.8049 | 0.6875 | 0.7416 | 300 |

Table 4: Performance of the Random Forest model with Random Forest Importance selection on different numbers of features.

| Model | SS | Accuracy | Precision | Recall | F1-score | # features |
|---|---|---|---|---|---|---|
| DT | Mutual Info | 0.7479 | 0.6800 | 0.7083 | 0.6939 | 50 |
| DT | Mutual Info | 0.7227 | 0.6471 | 0.6875 | 0.6667 | 60 |
| DT | Mutual Info | 0.7311 | 0.6600 | 0.6875 | 0.6735 | 70 |
| DT | Mutual Info | 0.7395 | 0.6889 | 0.6458 | 0.6667 | 80 |
| DT | Mutual Info | 0.7731 | 0.7561 | 0.6458 | 0.6966 | 90 |
| DT | Mutual Info | 0.7563 | 0.7209 | 0.6458 | 0.6813 | 100 |
| DT | Mutual Info | 0.7647 | 0.7273 | 0.6667 | 0.6957 | 110 |
| DT | Mutual Info | 0.7731 | 0.7333 | 0.6875 | 0.7097 | 120 |
| DT | Mutual Info | 0.7563 | 0.7209 | 0.6458 | 0.6813 | 130 |
| **DT** | **Mutual Info** | **0.8067** | **0.7551** | **0.7708** | **0.7629** | **140** |
| DT | Mutual Info | 0.7563 | 0.7021 | 0.6875 | 0.6947 | 150 |
| DT | Mutual Info | 0.7731 | 0.7333 | 0.6875 | 0.7097 | 160 |
| DT | Mutual Info | 0.7563 | 0.6863 | 0.7292 | 0.7071 | 170 |
| DT | Mutual Info | 0.7479 | 0.6957 | 0.6667 | 0.6809 | 180 |
| DT | Mutual Info | 0.7647 | 0.7000 | 0.7292 | 0.7143 | 190 |
| DT | Mutual Info | 0.7143 | 0.6296 | 0.7083 | 0.6667 | 200 |
| DT | Mutual Info | 0.7395 | 0.6977 | 0.6250 | 0.6593 | 210 |
| DT | Mutual Info | 0.7479 | 0.7045 | 0.6458 | 0.6739 | 220 |
| DT | Mutual Info | 0.7563 | 0.7021 | 0.6875 | 0.6947 | 230 |
| DT | Mutual Info | 0.7479 | 0.6875 | 0.6875 | 0.6875 | 240 |
| DT | Mutual Info | 0.7647 | 0.7174 | 0.6875 | 0.7021 | 250 |
| DT | Mutual Info | 0.7731 | 0.7143 | 0.7292 | 0.7216 | 260 |
| DT | Mutual Info | 0.7479 | 0.6800 | 0.7083 | 0.6939 | 270 |
| DT | Mutual Info | 0.7647 | 0.7273 | 0.6667 | 0.6957 | 280 |
| DT | Mutual Info | 0.7563 | 0.7317 | 0.6250 | 0.6742 | 290 |
| DT | Mutual Info | 0.7479 | 0.6957 | 0.6667 | 0.6809 | 300 |

Table 5: Performance of the Decision Tree model with Mutual Information selection on different numbers of features.

| Model | SS | Accuracy | Precision | Recall | F1-score | n features |
|---|---|---|---|---|---|---|
| DT | ANOVA | 0.7394 | 0.6666 | 0.7083 | 0.6868 | 50 |
| DT | ANOVA | 0.7563 | 0.6862 | 0.7291 | 0.7070 | 60 |
| DT | ANOVA | 0.7647 | 0.7173 | 0.6875 | 0.7021 | 70 |
| DT | ANOVA | 0.7647 | 0.7173 | 0.6875 | 0.7021 | 80 |
| DT | ANOVA | 0.7647 | 0.6851 | 0.7708 | 0.7254 | 90 |
| DT | ANOVA | 0.7815 | 0.7500 | 0.6875 | 0.7173 | 100 |
| DT | ANOVA | 0.7226 | 0.6595 | 0.6458 | 0.6526 | 110 |
| DT | ANOVA | 0.7563 | 0.7111 | 0.6666 | 0.6881 | 120 |
| DT | ANOVA | 0.7731 | 0.7333 | 0.6875 | 0.7096 | 130 |
| DT | ANOVA | 0.7899 | 0.7346 | 0.7500 | 0.7422 | 140 |
| **DT** | **ANOVA** | **0.7983** | **0.7608** | **0.7291** | **0.7446** | **150** |
| DT | ANOVA | 0.7731 | 0.7234 | 0.7083 | 0.7157 | 160 |
| DT | ANOVA | 0.7899 | 0.7173 | 0.6875 | 0.7021 | 170 |
| DT | ANOVA | 0.7983 | 0.7446 | 0.7291 | 0.7368 | 180 |
| DT | ANOVA | 0.7731 | 0.7234 | 0.7083 | 0.7157 | 190 |
| DT | ANOVA | 0.7563 | 0.7111 | 0.6666 | 0.6881 | 200 |
| DT | ANOVA | 0.6974 | 0.6363 | 0.5833 | 0.6086 | 210 |
| DT | ANOVA | 0.7058 | 0.6444 | 0.6041 | 0.6236 | 220 |
| DT | ANOVA | 0.7563 | 0.7021 | 0.6875 | 0.6947 | 230 |
| DT | ANOVA | 0.6974 | 0.7045 | 0.6458 | 0.6739 | 240 |
| DT | ANOVA | 0.6806 | 0.6041 | 0.6041 | 0.6041 | 250 |
| DT | ANOVA | 0.6806 | 0.6041 | 0.6041 | 0.6041 | 260 |
| DT | ANOVA | 0.7310 | 0.6818 | 0.6250 | 0.6521 | 270 |
| DT | ANOVA | 0.7226 | 0.6829 | 0.5833 | 0.6292 | 280 |
| DT | ANOVA | 0.7310 | 0.6904 | 0.6041 | 0.6444 | 290 |
| DT | ANOVA | 0.7226 | 0.6666 | 0.6250 | 0.6451 | 300 |

Table 6: Performance of the Decision Tree model with ANOVA selection on different numbers of features.

| **Model** | **SS** | **Accuracy** | **Precision** | **Recall** | **F1** | **# features** |
|---|---|---|---|---|---|---|
| DT | Chi2 | 0.7563 | 0.7436 | 0.6042 | 0.6667 | 50 |
| DT | Chi2 | 0.7899 | 0.7805 | 0.6667 | 0.7191 | 60 |
| DT | Chi2 | 0.7311 | 0.6600 | 0.6875 | 0.6735 | 70 |
| DT | Chi2 | 0.7647 | 0.7083 | 0.7083 | 0.7083 | 80 |
| DT | Chi2 | 0.7311 | 0.6600 | 0.6875 | 0.6735 | 90 |
| DT | Chi2 | 0.6639 | 0.5769 | 0.6250 | 0.6000 | 100 |
| DT | Chi2 | 0.6975 | 0.6071 | 0.7083 | 0.6538 | 110 |
| DT | Chi2 | 0.7395 | 0.6735 | 0.6875 | 0.6804 | 120 |
| DT | Chi2 | 0.7227 | 0.6531 | 0.6667 | 0.6598 | 130 |
| DT | Chi2 | 0.7479 | 0.6731 | 0.7292 | 0.7000 | 140 |
| DT | Chi2 | 0.7563 | 0.6862 | 0.7292 | 0.7070 | 150 |
| DT | Chi2 | 0.7478 | 0.6956 | 0.6666 | 0.6808 | 160 |
| DT | Chi2 | 0.7142 | 0.6346 | 0.6875 | 0.6600 | 170 |
| **DT** | **Chi2** | **0.7899** | **0.7555** | **0.7083** | **0.7311** | **180** |
| DT | Chi2 | 0.7394 | 0.6734 | 0.6875 | 0.6804 | 190 |
| DT | Chi2 | 0.7815 | 0.7391 | 0.7083 | 0.7234 | 200 |
| DT | Chi2 | 0.7563 | 0.6862 | 0.7291 | 0.7070 | 210 |
| DT | Chi2 | 0.7394 | 0.6888 | 0.6458 | 0.6666 | 220 |
| DT | Chi2 | 0.7394 | 0.6888 | 0.6458 | 0.6666 | 230 |
| DT | Chi2 | 0.7310 | 0.6739 | 0.6458 | 0.6595 | 240 |
| DT | Chi2 | 0.7478 | 0.6800 | 0.7083 | 0.6938 | 250 |
| DT | Chi2 | 0.7647 | 0.6923 | 0.7500 | 0.7200 | 260 |
| DT | Chi2 | 0.6974 | 0.6250 | 0.6250 | 0.6250 | 270 |
| DT | Chi2 | 0.7142 | 0.6400 | 0.6666 | 0.6530 | 280 |
| DT | Chi2 | 0.7142 | 0.6296 | 0.7083 | 0.6666 | 290 |
| DT | Chi2 | 0.7226 | 0.6595 | 0.6458 | 0.6526 | 300 |

Table 7: Performance of the Decision Tree model with Chi2 selection on different numbers of features.

| Model | SS | Accuracy | Precision | Recall | F1 | # features |
|---|---|---|---|---|---|---|
| NB | ANOVA | 0.6387 | 0.5510 | 0.5625 | 0.5567 | 60 |
| NB | ANOVA | 0.6134 | 0.5200 | 0.5417 | 0.5306 | 70 |
| NB | ANOVA | 0.6218 | 0.5306 | 0.5417 | 0.5361 | 80 |
| NB | ANOVA | 0.6303 | 0.5400 | 0.5625 | 0.5510 | 90 |
| NB | ANOVA | 0.6387 | 0.5510 | 0.5625 | 0.5567 | 100 |
| NB | ANOVA | 0.6303 | 0.5417 | 0.5417 | 0.5417 | 110 |
| NB | ANOVA | 0.6387 | 0.5532 | 0.5417 | 0.5474 | 120 |
| NB | ANOVA | 0.6303 | 0.5400 | 0.5625 | 0.5510 | 130 |
| NB | ANOVA | 0.6387 | 0.5510 | 0.5625 | 0.5567 | 140 |
| NB | ANOVA | 0.6303 | 0.5400 | 0.5625 | 0.5510 | 150 |
| NB | ANOVA | 0.6218 | 0.5294 | 0.5625 | 0.5455 | 160 |
| NB | ANOVA | 0.6218 | 0.5294 | 0.5625 | 0.5455 | 170 |
| NB | ANOVA | 0.6303 | 0.5385 | 0.5833 | 0.5600 | 180 |
| **NB** | **ANOVA** | **0.6387** | **0.5490** | **0.5833** | **0.5657** | **190** |
| **NB** | **ANOVA** | **0.6387** | **0.5490** | **0.5833** | **0.5657** | **200** |
| NB | ANOVA | 0.6303 | 0.5400 | 0.5625 | 0.5510 | 210 |
| NB | ANOVA | 0.6387 | 0.5510 | 0.5625 | 0.5567 | 220 |
| NB | ANOVA | 0.6387 | 0.5510 | 0.5625 | 0.5567 | 230 |
| NB | ANOVA | 0.6303 | 0.5400 | 0.5625 | 0.5510 | 240 |
| NB | ANOVA | 0.6471 | 0.5625 | 0.5625 | 0.5625 | 250 |
| NB | ANOVA | 0.6387 | 0.5510 | 0.5625 | 0.5567 | 260 |
| NB | ANOVA | 0.6303 | 0.5417 | 0.5417 | 0.5417 | 270 |
| NB | ANOVA | 0.6303 | 0.5400 | 0.5625 | 0.5510 | 280 |
| NB | ANOVA | 0.6303 | 0.5400 | 0.5625 | 0.5510 | 290 |
| NB | ANOVA | 0.6387 | 0.5510 | 0.5625 | 0.5567 | 300 |

Table 8: Performance of the Naive Bayes model with ANOVA selection on different numbers of features.

| Model | SS | Accuracy | Precision | Recall | F1 | # features |
|---|---|---|---|---|---|---|
| NB | Chi2 | 0.6975 | 0.7000 | 0.4375 | 0.5385 | 60 |
| NB | Chi2 | 0.6975 | 0.6875 | 0.4583 | 0.5500 | 70 |
| NB | Chi2 | 0.6723 | 0.6216 | 0.4792 | 0.5412 | 80 |
| NB | Chi2 | 0.6807 | 0.6389 | 0.4792 | 0.5476 | 90 |
| NB | Chi2 | 0.6639 | 0.6053 | 0.4792 | 0.5349 | 100 |
| NB | Chi2 | 0.6807 | 0.6316 | 0.5000 | 0.5581 | 110 |
| NB | Chi2 | 0.6891 | 0.6410 | 0.5208 | 0.5747 | 120 |
| NB | Chi2 | 0.7059 | 0.6667 | 0.5417 | 0.5977 | 130 |
| NB | Chi2 | 0.6723 | 0.5763 | 0.7083 | 0.6355 | 140 |
| NB | Chi2 | 0.6555 | 0.5507 | 0.7917 | 0.6496 | 150 |
| NB | Chi2 | 0.6555 | 0.5507 | 0.7917 | 0.6496 | 160 |
| NB | Chi2 | 0.6723 | 0.5672 | 0.7917 | 0.6609 | 170 |
| NB | Chi2 | 0.6387 | 0.5342 | 0.8125 | 0.6446 | 180 |
| NB | Chi2 | 0.6723 | 0.5672 | 0.7917 | 0.6609 | 190 |
| NB | Chi2 | 0.6807 | 0.5758 | 0.7917 | 0.6667 | 200 |
| **NB** | **Chi2** | **0.6891** | **0.5846** | **0.7917** | **0.6726** | **210** |
| NB | Chi2 | 0.6723 | 0.5714 | 0.7500 | 0.6486 | 220 |
| NB | Chi2 | 0.6639 | 0.5645 | 0.7292 | 0.6364 | 230 |
| NB | Chi2 | 0.6723 | 0.5672 | 0.7917 | 0.6609 | 240 |
| NB | Chi2 | 0.6387 | 0.5352 | 0.7917 | 0.6387 | 250 |
| NB | Chi2 | 0.6303 | 0.5286 | 0.7708 | 0.6271 | 260 |
| NB | Chi2 | 0.6387 | 0.5362 | 0.7708 | 0.6325 | 270 |
| NB | Chi2 | 0.6975 | 0.6200 | 0.6458 | 0.6327 | 280 |
| NB | Chi2 | 0.6975 | 0.6200 | 0.6458 | 0.6327 | 290 |
| NB | Chi2 | 0.6975 | 0.6200 | 0.6458 | 0.6327 | 300 |

Table 9: Performance of the Naive Bayes model with Chi2 selection on different numbers of features.

| Modello | Metodo selezione | Accuracy | Precision | Recall | F1-score | n_features |
|---|---|---|---|---|---|---|
| NB | Mutual Info | 0.6555 | 0.5660 | 0.6250 | 0.5941 | 60 |
| NB | Mutual Info | 0.6723 | 0.5957 | 0.5833 | 0.5895 | 70 |
| NB | Mutual Info | 0.6471 | 0.5556 | 0.6250 | 0.5882 | 80 |
| NB | Mutual Info | 0.6723 | 0.5818 | 0.6667 | 0.6214 | 90 |
| **NB** | **Mutual Info** | **0.6891** | **0.6000** | **0.6875** | **0.6408** | **100** |
| NB | Mutual Info | 0.6639 | 0.5741 | 0.6458 | 0.6078 | 110 |
| NB | Mutual Info | 0.6471 | 0.5536 | 0.6458 | 0.5962 | 120 |
| NB | Mutual Info | 0.6134 | 0.5161 | 0.6667 | 0.5818 | 130 |
| NB | Mutual Info | 0.6218 | 0.5263 | 0.6250 | 0.5714 | 140 |
| NB | Mutual Info | 0.6303 | 0.5357 | 0.6250 | 0.5769 | 150 |
| NB | Mutual Info | 0.6387 | 0.5424 | 0.6667 | 0.5981 | 160 |
| NB | Mutual Info | 0.6387 | 0.5439 | 0.6458 | 0.5905 | 170 |
| NB | Mutual Info | 0.6387 | 0.5455 | 0.6250 | 0.5825 | 180 |
| NB | Mutual Info | 0.6387 | 0.5472 | 0.6042 | 0.5743 | 190 |
| NB | Mutual Info | 0.6387 | 0.5490 | 0.5833 | 0.5657 | 200 |
| NB | Mutual Info | 0.6387 | 0.5439 | 0.6458 | 0.5905 | 210 |
| NB | Mutual Info | 0.6303 | 0.5370 | 0.6042 | 0.5686 | 220 |
| NB | Mutual Info | 0.6639 | 0.5741 | 0.6458 | 0.6078 | 230 |
| NB | Mutual Info | 0.6471 | 0.5536 | 0.6458 | 0.5962 | 240 |
| NB | Mutual Info | 0.6555 | 0.5686 | 0.6042 | 0.5859 | 250 |
| NB | Mutual Info | 0.6723 | 0.5882 | 0.6250 | 0.6061 | 260 |
| NB | Mutual Info | 0.6471 | 0.5600 | 0.5833 | 0.5714 | 270 |
| NB | Mutual Info | 0.6471 | 0.5714 | 0.5000 | 0.5333 | 280 |
| NB | Mutual Info | 0.6471 | 0.5625 | 0.5625 | 0.5625 | 290 |
| NB | Mutual Info | 0.6050 | 0.5000 | 0.5417 | 0.5253 | 300 |

Table 10: Performance of the Naive Bayes model with Mutual Info selection on different numbers of features.

### 2 Summary of Feature Selection using CROSS VALIDATION

#### F1-Score by Model, Feature Selection Method, and Number of Features

To evaluate model performance, we focus on the **F1-score**, as it best captures the balance between precision and recall. For each combination of model, feature selection method, and number of selected features, we report the **mean F1-score** across cross-validation folds. This allows a fair comparison of model effectiveness under different feature selection strategies and feature set sizes.

| Model | SS | # features | Accuracy | Precision | Recall | F1 |
|---|---|---|---|---|---|---|
| DecisionTree | ANOVA | 50 | 0.78616 | 0.7377 | 0.7272 | 0.7307 |
| DecisionTree | ANOVA | 60 | 0.7769 | 0.7245 | 0.7203 | 0.7191 |
| DecisionTree | ANOVA | 70 | 0.7760 | 0.7200 | 0.7209 | 0.7175 |
| DecisionTree | ANOVA | 80 | 0.7727 | 0.7082 | 0.7328 | 0.7186 |
| DecisionTree | ANOVA | 90 | 0.7735 | 0.7151 | 0.7230 | 0.7160 |

| | | | | | | |
|---|---|---|---|---|---|---|
| DecisionTree | ANOVA | 100 | 0.7744 | 0.7230 | 0.7185 | 0.7162 |
| DecisionTree | ANOVA | 110 | 0.7483 | 0.6892 | 0.6990 | 0.6878 |
| DecisionTree | ANOVA | 120 | 0.7558 | 0.6959 | 0.6921 | 0.6907 |
| DecisionTree | ANOVA | 130 | 0.7491 | 0.6907 | 0.6863 | 0.6843 |
| DecisionTree | ANOVA | 140 | 0.7642 | 0.7026 | 0.7002 | 0.6998 |
| DecisionTree | ANOVA | 150 | 0.7777 | 0.7145 | 0.7321 | 0.7216 |
| DecisionTree | ANOVA | 160 | 0.7802 | 0.7106 | 0.7472 | 0.7271 |
| DecisionTree | ANOVA | 170 | 0.7626 | 0.6972 | 0.7229 | 0.7075 |
| DecisionTree | ANOVA | 180 | 0.7592 | 0.6890 | 0.7262 | 0.7047 |
| DecisionTree | ANOVA | 190 | 0.7802 | 0.7107 | 0.7585 | 0.7323 |
| DecisionTree | ANOVA | 200 | 0.7735 | 0.7113 | 0.7393 | 0.7217 |
| DecisionTree | ANOVA | 210 | 0.7726 | 0.7081 | 0.7354 | 0.7192 |
| DecisionTree | ANOVA | 220 | 0.7962 | 0.7368 | 0.7646 | 0.7486 |
| DecisionTree | ANOVA | 230 | 0.7853 | 0.7290 | 0.7388 | 0.7322 |
| DecisionTree | ANOVA | 240 | 0.7937 | 0.7530 | 0.7238 | 0.7352 |
| DecisionTree | ANOVA | 250 | 0.7878 | 0.7346 | 0.7379 | 0.7336 |
| DecisionTree | ANOVA | 260 | 0.7735 | 0.7223 | 0.7163 | 0.7154 |
| DecisionTree | ANOVA | 270 | 0.7768 | 0.7194 | 0.7231 | 0.7192 |
| DecisionTree | ANOVA | 280 | 0.7769 | 0.7244 | 0.7267 | 0.7206 |
| DecisionTree | ANOVA | 290 | 0.7701 | 0.7189 | 0.7075 | 0.7096 |
| DecisionTree | ANOVA | 300 | 0.7659 | 0.7111 | 0.7159 | 0.7090 |
| DecisionTree | Chi2 | 50 | 0.7718 | 0.7088 | 0.7332 | 0.7177 |
| DecisionTree | Chi2 | 60 | 0.7811 | 0.7242 | 0.7388 | 0.7274 |
| DecisionTree | Chi2 | 70 | 0.7676 | 0.6977 | 0.7411 | 0.7165 |
| DecisionTree | Chi2 | 80 | 0.7718 | 0.7095 | 0.7304 | 0.7175 |
| DecisionTree | Chi2 | 90 | 0.7760 | 0.7278 | 0.7245 | 0.7194 |
| DecisionTree | Chi2 | 100 | 0.7643 | 0.7040 | 0.7088 | 0.7044 |
| DecisionTree | Chi2 | 110 | 0.7777 | 0.7182 | 0.7268 | 0.7204 |
| DecisionTree | Chi2 | 120 | 0.7837 | 0.7251 | 0.7471 | 0.7332 |
| DecisionTree | Chi2 | 130 | 0.7861 | 0.7216 | 0.7563 | 0.7368 |
| DecisionTree | Chi2 | 140 | 0.7793 | 0.7202 | 0.7336 | 0.7246 |
| DecisionTree | Chi2 | 150 | 0.7642 | 0.6995 | 0.7157 | 0.7058 |
| DecisionTree | Chi2 | 160 | 0.7743 | 0.7088 | 0.7416 | 0.7218 |
| DecisionTree | Chi2 | 170 | 0.7785 | 0.7117 | 0.7506 | 0.7272 |
| DecisionTree | Chi2 | 180 | 0.7769 | 0.7107 | 0.7449 | 0.7257 |
| DecisionTree | Chi2 | 190 | 0.7668 | 0.6997 | 0.7215 | 0.7086 |
| DecisionTree | Chi2 | 200 | 0.7777 | 0.7162 | 0.7353 | 0.7228 |
| DecisionTree | Chi2 | 210 | 0.7626 | 0.6947 | 0.7264 | 0.7079 |
| DecisionTree | Chi2 | 220 | 0.7584 | 0.6921 | 0.7082 | 0.6986 |
| DecisionTree | Chi2 | 230 | 0.7542 | 0.6834 | 0.7186 | 0.6986 |
| DecisionTree | Chi2 | 240 | 0.7660 | 0.7065 | 0.7122 | 0.7064 |
| DecisionTree | Chi2 | 250 | 0.7559 | 0.6833 | 0.7258 | 0.7013 |
| DecisionTree | Chi2 | 260 | 0.7642 | 0.6997 | 0.7234 | 0.7092 |
| DecisionTree | Chi2 | 270 | 0.7634 | 0.6998 | 0.7245 | 0.7086 |
| DecisionTree | Chi2 | 280 | 0.7710 | 0.7089 | 0.7340 | 0.7188 |
| DecisionTree | Chi2 | 290 | 0.7727 | 0.7084 | 0.7391 | 0.7192 |

| | | | | | | |
|---|---|---|---|---|---|---|
| DecisionTree | Chi2 | 300 | 0.7592 | 0.6859 | 0.7355 | 0.7071 |
| DecisionTree | MutualInfo | 50 | 0.7929 | 0.7489 | 0.7339 | 0.7383 |
| DecisionTree | MutualInfo | 60 | 0.7879 | 0.7373 | 0.7384 | 0.7346 |
| DecisionTree | MutualInfo | 70 | 0.7920 | 0.7360 | 0.7493<br>9 | 0.7409 |
| DecisionTree | MutualInfo | 80 | 0.79463 | 0.7456 | 0.7450 | 0.7427 |
| DecisionTree | MutualInfo | 90 | 0.7988 | 0.7504 | 0.7420 | 0.7446 |
| DecisionTree | MutualInfo | 100 | 0.7903 | 0.7404 | 0.7333 | 0.7340 |
| DecisionTree | MutualInfo | 110 | 0.7786 | 0.7190 | 0.7390 | 0.7270 |
| DecisionTree | MutualInfo | 120 | 0.7971 | 0.7529 | 0.7428 | 0.7453 |
| DecisionTree | MutualInfo | 130 | 0.7938 | 0.7317 | 0.7624 | 0.7461 |
| DecisionTree | MutualInfo | 140 | 0.7895 | 0.7362 | 0.7423 | 0.7372 |
| DecisionTree | MutualInfo | 150 | 0.8005 | 0.7459 | 0.7634 | 0.7527 |
| DecisionTree | MutualInfo | 160 | 0.7752 | 0.7087 | 0.7479 | 0.7258 |
| DecisionTree | MutualInfo | 170 | 0.7920 | 0.7331 | 0.7598 | 0.7445 |
| DecisionTree | MutualInfo | 180 | 0.7853 | 0.7212 | 0.7519 | 0.7350 |
| DecisionTree | MutualInfo | 190 | 0.7837 | 0.7195 | 0.7606 | 0.7383 |
| DecisionTree | MutualInfo | 200 | 0.7996 | 0.7394 | 0.7727 | 0.7544 |
| DecisionTree | MutualInfo | 210 | 0.7828 | 0.7240 | 0.7431 | 0.7314 |
| DecisionTree | MutualInfo | 220 | 0.7811 | 0.7179 | 0.7532 | 0.7332 |
| DecisionTree | MutualInfo | 230 | 0.7677 | 0.6983 | 0.7380 | 0.7157 |
| DecisionTree | MutualInfo | 240 | 0.7634 | 0.6968 | 0.7213 | 0.7076 |
| DecisionTree | MutualInfo | 250 | 0.7710 | 0.7068 | 0.7461 | 0.7222 |
| DecisionTree | MutualInfo | 260 | 0.7903 | 0.7272 | 0.7571 | 0.7405 |
| DecisionTree | MutualInfo | 270 | 0.7769 | 0.7087 | 0.7573 | 0.7305 |
| DecisionTree | MutualInfo | 280 | 0.7719 | 0.7160 | 0.7169 | 0.7146 |
| DecisionTree | MutualInfo | 290 | 0.7592 | 0.6921 | 0.7286 | 0.7067 |
| DecisionTree | MutualInfo | 300 | 0.7701 | 0.6974 | 0.7376 | 0.7163 |

| Model | SS | Num Features | Accuracy | Precision | Recall | F1 |
|---|---|---|---|---|---|---|
| NaiveBayes | ANOVA | 50 | 0.6842 | 0.6063 | 0.6241 | 0.6091 |
| NaiveBayes | ANOVA | 60 | 0.6893 | 0.6098 | 0.6416 | 0.6202 |
| NaiveBayes | ANOVA | 70 | 0.6842 | 0.6020 | 0.6416 | 0.6163 |
| NaiveBayes | ANOVA | 80 | 0.6868 | 0.6055 | 0.6404 | 0.6173 |
| NaiveBayes | ANOVA | 90 | 0.6809 | 0.5983 | 0.6407 | 0.6129 |
| NaiveBayes | ANOVA | 100 | 0.6809 | 0.5987 | 0.6427 | 0.6143 |
| NaiveBayes | ANOVA | 110 | 0.6809 | 0.6012 | 0.6270 | 0.6084 |
| NaiveBayes | ANOVA | 120 | 0.6842 | 0.6036 | 0.6384 | 0.6150 |
| NaiveBayes | ANOVA | 130 | 0.6851 | 0.6052 | 0.6362 | 0.6148 |
| NaiveBayes | ANOVA | 140 | 0.6885 | 0.6069 | 0.6458 | 0.6210 |
| NaiveBayes | ANOVA | 150 | 0.6918 | 0.6111 | 0.6460 | 0.6230 |
| NaiveBayes | ANOVA | 160 | 0.6851 | 0.6021 | 0.6414 | 0.6164 |
| NaiveBayes | ANOVA | 170 | 0.6851 | 0.6010 | 0.6463 | 0.6176 |
| NaiveBayes | ANOVA | 180 | 0.6851 | 0.6019 | 0.6422 | 0.6160 |

| | | | | | | |
|---|---|---|---|---|---|---|
| NaiveBayes | ANOVA | 190 | 0.6876 | 0.6063 | 0.6447 | 0.6197 |
| NaiveBayes | ANOVA | 200 | 0.6868 | 0.6049 | 0.6449 | 0.6186 |
| NaiveBayes | ANOVA | 210 | 0.6876 | 0.6056 | 0.6446 | 0.6195 |
| NaiveBayes | ANOVA | 220 | 0.6860 | 0.6038 | 0.6446 | 0.6183 |
| NaiveBayes | ANOVA | 230 | 0.6851 | 0.6046 | 0.6361 | 0.6144 |
| NaiveBayes | ANOVA | 240 | 0.6877 | 0.6072 | 0.6383 | 0.6168 |
| NaiveBayes | ANOVA | 250 | 0.6902 | 0.6110 | 0.6385 | 0.6189 |
| NaiveBayes | ANOVA | 260 | 0.6876 | 0.6077 | 0.6342 | 0.6149 |
| NaiveBayes | ANOVA | 270 | 0.6885 | 0.6098 | 0.6299 | 0.6142 |
| NaiveBayes | ANOVA | 280 | 0.6851 | 0.6059 | 0.6184 | 0.6067 |
| NaiveBayes | ANOVA | 290 | 0.6834 | 0.6056 | 0.6122 | 0.6038 |
| NaiveBayes | ANOVA | 300 | 0.6826 | 0.6029 | 0.6160 | 0.6044 |
| NaiveBayes | Chi2 | 50 | 0.6818 | 0.6588 | 0.4245 | 0.5110 |
| NaiveBayes | Chi2 | 60 | 0.6784 | 0.6541 | 0.4488 | 0.5223 |
| NaiveBayes | Chi2 | 70 | 0.6910 | 0.6582 | 0.4985 | 0.5583 |
| NaiveBayes | Chi2 | 80 | 0.6893 | 0.6483 | 0.5050 | 0.5518 |
| NaiveBayes | Chi2 | 90 | 0.6910 | 0.6363 | 0.5265 | 0.5661 |
| NaiveBayes | Chi2 | 100 | 0.6969 | 0.6422 | 0.5560 | 0.5906 |
| NaiveBayes | Chi2 | 110 | 0.7020 | 0.6504 | 0.5805 | 0.6001 |
| NaiveBayes | Chi2 | 120 | 0.7003 | 0.6304 | 0.6252 | 0.6164 |
| NaiveBayes | Chi2 | 130 | 0.6809 | 0.6027 | 0.6576 | 0.6129 |
| NaiveBayes | Chi2 | 140 | 0.6650 | 0.5825 | 0.6742 | 0.6110 |
| NaiveBayes | Chi2 | 150 | 0.6624 | 0.5757 | 0.6836 | 0.6130 |
| NaiveBayes | Chi2 | 160 | 0.6583 | 0.5651 | 0.7299 | 0.6250 |
| NaiveBayes | Chi2 | 170 | 0.6575 | 0.5644 | 0.7469 | 0.6322 |
| NaiveBayes | Chi2 | 180 | 0.6608 | 0.5661 | 0.7525 | 0.6364 |
| NaiveBayes | Chi2 | 190 | 0.6499 | 0.5489 | 0.7565 | 0.6302 |
| NaiveBayes | Chi2 | 200 | 0.6490 | 0.5477 | 0.7754 | 0.6368 |
| NaiveBayes | Chi2 | 210 | 0.6440 | 0.5400 | 0.7980 | 0.6390 |
| NaiveBayes | Chi2 | 220 | 0.6330 | 0.5353 | 0.8050 | 0.6355 |
| NaiveBayes | Chi2 | 230 | 0.6297 | 0.5288 | 0.8210 | 0.6383 |
| NaiveBayes | Chi2 | 240 | 0.6330 | 0.5327 | 0.8228 | 0.6407 |
| NaiveBayes | Chi2 | 250 | 0.6330 | 0.5328 | 0.8204 | 0.6404 |
| NaiveBayes | Chi2 | 260 | 0.6313 | 0.5310 | 0.8121 | 0.6363 |
| NaiveBayes | Chi2 | 270 | 0.6423 | 0.5398 | 0.8062 | 0.6417 |
| NaiveBayes | Chi2 | 280 | 0.6153 | 0.5226 | 0.8372 | 0.6343 |
| NaiveBayes | Chi2 | 290 | 0.6170 | 0.5199 | 0.8572 | 0.6420 |
| NaiveBayes | Chi2 | 300 | 0.6144 | 0.5166 | 0.8581 | 0.6400 |
| NaiveBayes | MutualInfo | 50 | 0.6809 | 0.5945 | 0.6504 | 0.6162 |
| NaiveBayes | MutualInfo | 60 | 0.6725 | 0.5868 | 0.6283 | 0.6019 |
| NaiveBayes | MutualInfo | 70 | 0.6817 | 0.6001 | 0.6378 | 0.6123 |
| NaiveBayes | MutualInfo | 80 | 0.6759 | 0.5956 | 0.6145 | 0.5990 |
| NaiveBayes | MutualInfo | 90 | 0.6759 | 0.6024 | 0.5830 | 0.5849 |
| NaiveBayes | MutualInfo | 100 | 0.6817 | 0.6010 | 0.6040 | 0.5995 |
| NaiveBayes | MutualInfo | 110 | 0.6834 | 0.6058 | 0.6110 | 0.6026 |
| NaiveBayes | MutualInfo | 120 | 0.6834 | 0.6036 | 0.6046 | 0.6002 |

| | | | | | | |
|---|---|---|---|---|---|---|
| NaiveBayes | MutualInfo | 130 | 0.6792 | 0.5985 | 0.6468 | 0.6131 |
| NaiveBayes | MutualInfo | 140 | 0.6759 | 0.5908 | 0.6528 | 0.6135 |
| NaiveBayes | MutualInfo | 150 | 0.6759 | 0.5931 | 0.6350 | 0.6063 |
| NaiveBayes | MutualInfo | 160 | 0.6826 | 0.5984 | 0.6549 | 0.6196 |
| NaiveBayes | MutualInfo | 170 | 0.6750 | 0.5905 | 0.6470 | 0.6104 |
| NaiveBayes | MutualInfo | 180 | 0.6759 | 0.5912 | 0.6382 | 0.6072 |
| NaiveBayes | MutualInfo | 190 | 0.6758 | 0.5919 | 0.6353 | 0.6063 |
| NaiveBayes | MutualInfo | 200 | 0.6759 | 0.5947 | 0.6275 | 0.6040 |
| NaiveBayes | MutualInfo | 210 | 0.6725 | 0.5918 | 0.6192 | 0.5981 |
| NaiveBayes | MutualInfo | 220 | 0.6759 | 0.5929 | 0.6297 | 0.6050 |
| NaiveBayes | MutualInfo | 230 | 0.6750 | 0.5921 | 0.6155 | 0.5975 |
| NaiveBayes | MutualInfo | 240 | 0.6742 | 0.5904 | 0.6184 | 0.5990 |
| NaiveBayes | MutualInfo | 250 | 0.6708 | 0.5883 | 0.6091 | 0.5930 |
| NaiveBayes | MutualInfo | 260 | 0.6767 | 0.5970 | 0.6072 | 0.5964 |
| NaiveBayes | MutualInfo | 270 | 0.6809 | 0.6007 | 0.6165 | 0.6037 |
| NaiveBayes | MutualInfo | 280 | 0.6750 | 0.5939 | 0.6155 | 0.5994 |
| NaiveBayes | MutualInfo | 290 | 0.6750 | 0.5954 | 0.6123 | 0.5972 |
| NaiveBayes | MutualInfo | 300 | 0.6784 | 0.5976 | 0.6171 | 0.6024 |

| Model | SS | Num Features | Accuracy | Precision | Recall | F1 |
|---|---|---|---|---|---|---|
| RandomForest | ANOVA | 50 | 0.8459 | 0.8212 | 0.7849 | 0.8007 |
| RandomForest | ANOVA | 60 | 0.8442 | 0.8142 | 0.7950 | 0.8013 |
| RandomForest | ANOVA | 70 | 0.8467 | 0.8216 | 0.7913 | 0.8036 |
| RandomForest | ANOVA | 80 | 0.8509 | 0.8255 | 0.7970 | 0.8087 |
| RandomForest | ANOVA | 90 | 0.8518 | 0.8252 | 0.7979 | 0.8097 |
| RandomForest | ANOVA | 100 | 0.8434 | 0.8196 | 0.7804 | 0.7974 |
| RandomForest | ANOVA | 110 | 0.8442 | 0.8116 | 0.7978 | 0.8018 |
| RandomForest | ANOVA | 120 | 0.8451 | 0.8199 | 0.7865 | 0.8002 |
| RandomForest | ANOVA | 130 | 0.8509 | 0.8261 | 0.7977 | 0.8093 |
| RandomForest | ANOVA | 140 | 0.8510 | 0.8248 | 0.7999 | 0.8089 |
| RandomForest | ANOVA | 150 | 0.8543 | 0.8309 | 0.8011 | 0.8130 |
| RandomForest | ANOVA | 160 | 0.8484 | 0.8233 | 0.7893 | 0.8040 |
| RandomForest | ANOVA | 170 | 0.8568 | 0.8321 | 0.8069 | 0.8168 |
| RandomForest | ANOVA | 180 | 0.8518 | 0.8325 | 0.7914 | 0.8083 |
| RandomForest | ANOVA | 190 | 0.8451 | 0.8185 | 0.78621 | 0.8003 |
| RandomForest | ANOVA | 200 | 0.8468 | 0.8248 | 0.7844 | 0.8019 |
| RandomForest | ANOVA | 210 | 0.8459 | 0.8184 | 0.7896 | 0.8013 |
| RandomForest | ANOVA | 220 | 0.8551 | 0.8365 | 0.7951 | 0.8128 |
| RandomForest | ANOVA | 230 | 0.8484 | 0.8198 | 0.7967 | 0.8058 |
| RandomForest | ANOVA | 240 | 0.8417 | 0.8216 | 0.7700 | 0.7926 |
| RandomForest | ANOVA | 250 | 0.8468 | 0.8289 | 0.7779 | 0.8001 |
| RandomForest | ANOVA | 260 | 0.8560 | 0.8353 | 0.8004 | 0.8148 |
| RandomForest | ANOVA | 270 | 0.8535 | 0.8310 | 0.7968 | 0.8114 |

| | | | | | | |
|---|---|---|---|---|---|---|
| RandomForest | ANOVA | 280 | 0.8467 | 0.8224 | 0.7886 | 0.8024 |
| RandomForest | ANOVA | 290 | 0.8585 | 0.8473 | 0.7907 | 0.8150 |
| RandomForest | ANOVA | 300 | 0.8467 | 0.8257 | 0.7832 | 0.8017 |
| RandomForest | Chi2 | 50 | 0.8602 | 0.8420 | 0.8013 | 0.8193 |
| RandomForest | Chi2 | 60 | 0.8610 | 0.8399 | 0.8043 | 0.8203 |
| RandomForest | Chi2 | 70 | 0.8501 | 0.8316 | 0.7882 | 0.8070 |
| RandomForest | Chi2 | 80 | 0.8560 | 0.8371 | 0.7996 | 0.8148 |
| RandomForest | Chi2 | 90 | 0.8543 | 0.8318 | 0.7993 | 0.8132 |
| RandomForest | Chi2 | 100 | 0.8551 | 0.8342 | 0.80138 | 0.8149 |
| RandomForest | Chi2 | 110 | 0.8585 | 0.8393 | 0.8030 | 0.8182 |
| RandomForest | Chi2 | 120 | 0.8518 | 0.8243 | 0.8019 | 0.8106 |
| RandomForest | Chi2 | 130 | 0.8577 | 0.8355 | 0.8034 | 0.8174 |
| RandomForest | Chi2 | 140 | 0.8560 | 0.8390 | 0.7941 | 0.8137 |
| RandomForest | Chi2 | 150 | 0.8543 | 0.8326 | 0.7996 | 0.8131 |
| RandomForest | Chi2 | 160 | 0.8476 | 0.8257 | 0.7841 | 0.8020 |
| RandomForest | Chi2 | 170 | 0.8534 | 0.8353 | 0.7901 | 0.8097 |
| RandomForest | Chi2 | 180 | 0.8493 | 0.8263 | 0.7905 | 0.8054 |
| RandomForest | Chi2 | 190 | 0.8526 | 0.8360 | 0.7871 | 0.8091 |
| RandomForest | Chi2 | 200 | 0.8535 | 0.8268 | 0.8043 | 0.8129 |
| RandomForest | Chi2 | 210 | 0.8568 | 0.8367 | 0.8035 | 0.8166 |
| RandomForest | Chi2 | 220 | 0.8526 | 0.8308 | 0.7959 | 0.8107 |
| RandomForest | Chi2 | 230 | 0.8551 | 0.8349 | 0.7973 | 0.8133 |
| RandomForest | Chi2 | 240 | 0.8543 | 0.8338 | 0.7950 | 0.8119 |
| RandomForest | Chi2 | 250 | 0.8450 | 0.8254 | 0.7829 | 0.8000 |
| RandomForest | Chi2 | 260 | 0.8535 | 0.8368 | 0.7892 | 0.8099 |
| RandomForest | Chi2 | 270 | 0.8560 | 0.8417 | 0.7893 | 0.8123 |
| RandomForest | Chi2 | 280 | 0.8568 | 0.8375 | 0.7972 | 0.8146 |
| RandomForest | Chi2 | 290 | 0.8560 | 0.8374 | 0.7989 | 0.8144 |
| RandomForest | Chi2 | 300 | 0.8543 | 0.8384 | 0.7916 | 0.8112 |
| RandomForest | MutualInfo | 50 | 0.8484 | 0.8258 | 0.7894 | 0.8044 |
| RandomForest | MutualInfo | 60 | 0.8459 | 0.8219 | 0.7903 | 0.8025 |
| RandomForest | MutualInfo | 70 | 0.8467 | 0.8199 | 0.7928 | 0.8034 |
| RandomForest | MutualInfo | 80 | 0.8501 | 0.8345 | 0.78476 | 0.8055 |
| RandomForest | MutualInfo | 90 | 0.8577 | 0.8405 | 0.7987 | 0.8164 |
| RandomForest | MutualInfo | 100 | 0.8568 | 0.8370 | 0.8017 | 0.8164 |
| RandomForest | MutualInfo | 110 | 0.8534 | 0.8359 | 0.7933 | 0.8109 |
| RandomForest | MutualInfo | 120 | 0.8526 | 0.8390 | 0.7845 | 0.8084 |
| RandomForest | MutualInfo | 130 | 0.8510 | 0.8298 | 0.7938 | 0.8085 |
| RandomForest | MutualInfo | 140 | 0.8526 | 0.8361 | 0.7918 | 0.8101 |
| RandomForest | MutualInfo | 150 | 0.8585 | 0.8439 | 0.7943 | 0.8158 |
| RandomForest | MutualInfo | 160 | 0.8518 | 0.8312 | 0.7899 | 0.8079 |
| RandomForest | MutualInfo | 170 | 0.8509 | 0.8330 | 0.7887 | 0.8074 |
| RandomForest | MutualInfo | 180 | 0.8552 | 0.8375 | 0.7963 | 0.8131 |
| RandomForest | MutualInfo | 190 | 0.8577 | 0.8358 | 0.8025 | 0.8165 |
| RandomForest | MutualInfo | 200 | 0.8510 | 0.8276 | 0.7939 | 0.8079 |
| RandomForest | MutualInfo | 210 | 0.8509 | 0.8269 | 0.7949 | 0.8082 |

| | | | | | | |
|---|---|---|---|---|---|---|
| RandomForest | MutualInfo | 220 | 0.8535 | 0.8351 | 0.7894 | 0.8095 |
| RandomForest | MutualInfo | 230 | 0.8585 | 0.8520 | 0.7840 | 0.8138 |
| RandomForest | MutualInfo | 240 | 0.8476 | 0.8267 | 0.7877 | 0.8042<br>3 |
| RandomForest | MutualInfo | 250 | 0.8518 | 0.8263 | 0.7993 | 0.8100 |
| RandomForest | MutualInfo | 260 | 0.8552 | 0.8337 | 0.7961 | 0.8126 |
| RandomForest | MutualInfo | 270 | 0.8611 | 0.8431 | 0.8060 | 0.8213 |
| RandomForest | MutualInfo | 280 | 0.8560 | 0.8390 | 0.7962 | 0.8145 |
| RandomForest | MutualInfo | 290 | 0.8619 | 0.8350 | 0.8156 | 0.8234 |
| RandomForest | MutualInfo | 300 | 0.8636 | 0.8431 | 0.8081 | 0.8237 |
| RandomForest | RFI | 50 | 0.8585 | 0.8324 | 0.8129 | 0.8198 |
| RandomForest | RFI | 60 | 0.8611 | 0.8396 | 0.8079 | 0.8213 |
| RandomForest | RFI | 70 | 0.8560 | 0.8308 | 0.8057 | 0.8152 |
| RandomForest | RFI | 80 | 0.8552 | 0.8346 | 0.7978 | 0.8135 |
| RandomForest | RFI | 90 | 0.8468 | 0.8223 | 0.7888 | 0.8023 |
| RandomForest | RFI | 100 | 0.8636 | 0.8470 | 0.8054 | 0.8235 |
| RandomForest | RFI | 110 | 0.8568 | 0.8303 | 0.8064 | 0.8163 |
| RandomForest | RFI | 120 | 0.8627 | 0.8484 | 0.8012 | 0.8217 |
| RandomForest | RFI | 130 | 0.8602 | 0.8461 | 0.7940 | 0.8172 |
| RandomForest | RFI | 140 | 0.8619 | 0.8475 | 0.8024 | 0.8211 |
| RandomForest | RFI | 150 | 0.8518 | 0.8347 | 0.7893 | 0.8078 |
| RandomForest | RFI | 160 | 0.8577 | 0.8420 | 0.7949 | 0.8154 |
| RandomForest | RFI | 170 | 0.8543 | 0.8379 | 0.7902 | 0.8103 |
| RandomForest | RFI | 180 | 0.8535 | 0.8397 | 0.7835 | 0.8086 |
| RandomForest | RFI | 190 | 0.8543 | 0.8344 | 0.7943 | 0.8114 |
| RandomForest | RFI | 200 | 0.8594 | 0.8416 | 0.7987 | 0.8172 |
| RandomForest | RFI | 210 | 0.8577 | 0.8461 | 0.7884 | 0.8132 |
| RandomForest | RFI | 220 | 0.8476 | 0.8252 | 0.7856 | 0.8025 |
| RandomForest | RFI | 230 | 0.8535 | 0.8341 | 0.7960 | 0.8112 |
| RandomForest | RFI | 240 | 0.8459 | 0.8340 | 0.7737 | 0.7985 |
| RandomForest | RFI | 250 | 0.8501 | 0.8292 | 0.7880 | 0.8060 |
| RandomForest | RFI | 260 | 0.8509 | 0.8267 | 0.7967 | 0.8083 |
| RandomForest | RFI | 270 | 0.8543 | 0.8357 | 0.7959 | 0.8118 |
| RandomForest | RFI | 280 | 0.8509 | 0.8348 | 0.7842 | 0.8060 |
| RandomForest | RFI | 290 | 0.8527 | 0.8431 | 0.7789 | 0.8067 |
| RandomForest | RFI | 300 | 0.8467 | 0.8235 | 0.7859 | 0.8017 |
